# Entropy of a bacterial stress response is a generalizable predictor for fitness and antibiotic sensitivity

**DOI:** 10.1101/813709

**Authors:** Zeyu Zhu, Defne Surujon, Juan C. Ortiz-Marquez, Stephen J. Wood, Wenwen Huo, Ralph R. Isberg, José Bento, Tim van Opijnen

**Affiliations:** Biology Department, Boston College, Chestnut Hill, MA, USA; Computer Science Department, Boston College, Chestnut Hill, MA, USA; Department of Molecular Biology and Microbiology, Tufts University School of Medicine, Boston, MA, USA

**Author notes:** Equal contribution (Zeyu Zhu, Defne Surujon).

**Keywords:** RNA-Seq, Tn-Seq, antibiotic resistance, systems biology, machine learning, data integration, predictive modeling

## Abstract

Genes implicated in bacterial stress responses have been used to construct models that infer the growth outcome of a bacterium in the presence of antibiotics with the objective to develop novel diagnostic methods in the clinic. Current models are trained on data specific to a species or type of stress, making them potentially limited in their application. It is unclear if a generalizable response-signature exists that can predict bacterial fitness independent of strain, species or type of stress. Here we present a substantial RNA-Seq and experimental evolution dataset for 9 strains and species, under multiple antibiotic and non-antibiotic stress conditions. We show that gene panel-based models can accurately predict antibiotic mechanism of action, as well as the fitness outcome of *Streptococcus pneumoniae* in the presence of antibiotics or under nutrient depletion. However, these models quickly become species-specific as gene homology is limited. Instead, we define a new concept, transcriptomic entropy, which we use to quantify the amount of transcriptional disruption that occurs in a bacterium when responding to the environment. With entropy at the center, we train a suite of predictive (machine learning) models enabling generalizable fitness and antibiotic sensitivity predictions. These entropy-based models that predict bacterial fitness are validated for 7 Gram-positive and -negative species under antibiotic and non-antibiotic conditions indicating that transcriptomic entropy can be used as a generalizable stress signature. Moreover, rather than being a binary indicator of fitness, an entropy-based model was developed and validated to predict the minimum inhibitory concentration of an antibiotic. Lastly, we show that the inclusion of a varied-set of multi-omics features of a bacterial stress response further enhances fitness predictions by reducing ambiguity. By demonstrating the feasibility of generalizable predictions of bacterial fitness, this work establishes the fundamentals for potentially new approaches in infectious disease diagnostics, including antibiotic susceptibility testing.

**Significance statement:** Accurate predictions of bacterial fitness outcome could potentially have clinical diagnostic value, such as predicting optimum antibiotic choice and dosage for treating infectious diseases. Existing models of fitness predictions rely mainly on gene panel approaches, which may be species- and stress-specific due to a lack of gene and response conservation. In order to overcome this limitation, we generated a substantial experimental dataset and identified entropy as a universal stress response signature that quantifies the level of transcriptional disruption that is indicative of fitness outcome under a stressful condition. We present and validate for Gram-positive and negative species a suite of entropy-based models that enable accurate predictions of fitness outcome and the level of antibiotic sensitivity in a species and stress-type independent manner.

## Introduction

It is generally assumed that in order to overcome a stress, bacteria activate a response such as the stringent response under nutrient deprivation (1) or the SOS response in the presence of DNA damage (2). Measuring the activation of a specific response, or genes associated with this response, can thereby function as an indicator of what type of stress is occurring in a bacterium. For instance, *lexA*, encoding a master regulator of the SOS response in *Escherichia coli* and *Salmonella* (3, 4) is upregulated in response to fluoroquinolones, indicative of the DNA damage resulting from this class of antibiotics (4). Moreover, genes implicated in a stress response can help construct statistical models for predicting growth outcomes under that stress. For instance, gene panels have been assembled from transcriptomic data to predict whether *E. coli* can successfully grow in the presence of antibiotics such as ciprofloxacin (5-7).

While a defined stress-response or a gene-panel can be valuable in identifying the type or sensitivity to a stress, there are several complications that make one-to-one implementation across strains, species or environments difficult. For instance, responses such as the stringent or SOS response are only well defined in a small number of species, genes in a gene-panel may not be conserved across strains or species, and responses are not necessarily regulated in the same manner in different strains or species (8, 9). This lack of well-characterized and/or conserved responses limits the potential generalizability of existing models that attempt to predict stress sensitivity across strains and species. Therefore, identification of a universal stress response signature would allow for the development of generalizable predictive models that work for any species under any condition. There is not, however, a generally agreed upon stress response signature.

We previously showed that a stress response can be captured on at least two organizational levels; with RNA-Seq genome-wide transcriptional changes upon an environmental perturbation can be described, while transposon-insertion sequencing (Tn-Seq) characterizes the phenotypic importance of a gene, i.e. a gene’s contribution to fitness in a specific environment, on a genome-wide scale (16-21). Direct comparisons of data obtained from these approaches revealed that genes that change in transcription are poor indicators of what matters phenotypically (16). In other words, phenotypically important and transcriptionally important genes rarely overlap (16). However, when integrated into a network, coordinated patterns between these data-sets surface when the organism is challenged with an evolutionarily familiar stress (i.e. one that has been experienced for many generations), while the response becomes less coordinated when the bacterium is challenged with and responds to a relatively new stress, for instance antibiotics (16). This suggests that a stress signature may exist that is indicative of the degree to which a bacterium is adapted to a specific stress and is able to overcome the challenge. To determine whether such a signature exists and whether it is universal we generated and mined a substantial experimental dataset for the bacterial pathogen *Streptococcus pneumoniae*. We first demonstrate that both an antibiotic’s MOA and the bacterium’s fitness under antibiotic or nutrient pressure can be predicted by expression profiles from two distinct small gene-panels. However, these panels are limited to strains and species sharing high sequence and response homology. Instead, we develop the concept of entropy and build a suite of predictive models that accurately predict a bacterium’s fitness outcome under various nutrient, antibiotic and transcriptional perturbation conditions. Moreover, entropy is quantitatively correlated with the level of antibiotic sensitivity, enabling MIC predictions. We show the universality of entropy by validation experiments with 7 Gram-negative and -positive pathogenic species. We further demonstrate that entropy-based models can be improved in accuracy when different data-types are combined into a Support Vector Machine. Overall, we develop a large new experimental dataset, a novel species-independent stress-response concept and a suite of predictive models that accommodate different amounts and types of data to enable generalizable fitness predictions and antibiotic sensitivity.

## Results

### Antibiotic transcriptional responses are distinguishable based on mechanisms of action

To determine if the transcriptional responses to antibiotics were dependent on their mechanisms of action, the TIGR4 (T4) and Taiwan-19F (19F) strains of *Streptococcus pneumoniae* were grown in the presence or absence of 1x the minimum inhibitory concentration (MIC) of 9 antibiotics representing 4 mechanisms of action (MOA’s). These include cell wall synthesis inhibitors (CWSI), DNA synthesis inhibitors (DSI), protein synthesis inhibitors (PSI) and RNA synthesis inhibitors ((RSI); **Figure 1A**, Supplemental Tables 1, 6). Each strain was exposed to each antibiotic for 2 to 4 hours and cells were harvested for RNA-Seq at various time points after antibiotic exposure. Previous models have predicted bacterial fitness of *Escherichia coli* exposed to specific stressful conditions by relying on differential expression of selected small gene sets (5-7). Before we set out to determine whether a generalizable feature can predict fitness across a variety of stress types, we first established if a small gene set can predict the MOA of an antibiotic and/or fitness of *S. pneumoniae* (**Figure 1A**). Principal component analysis (PCA) was performed on the complete differential transcription datasets, and this indicated that temporal transcriptional responses to drugs within the same MOA tended to follow similar trajectories over time (**Figure 1B, C**).

**Figure 1.**
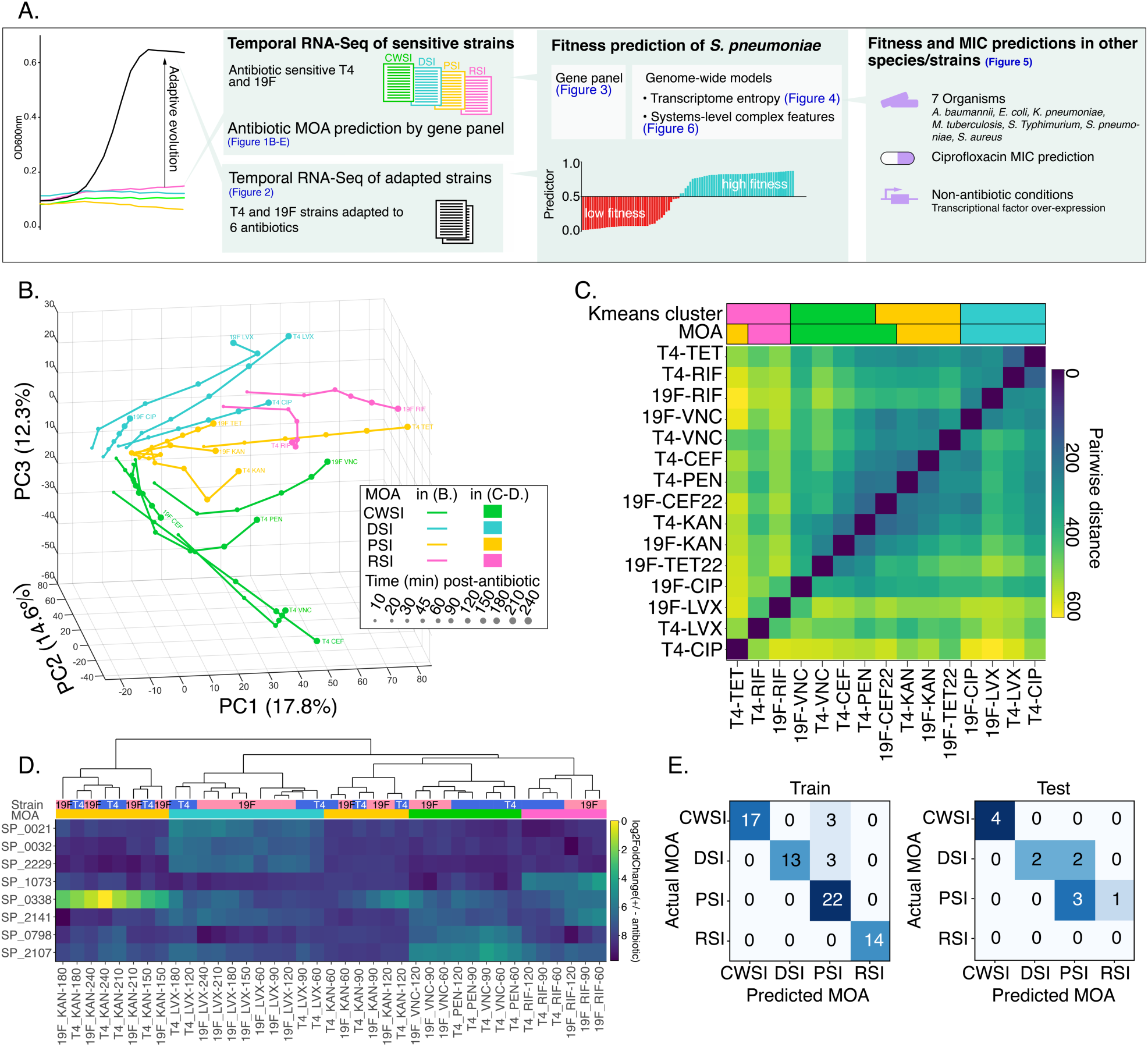
Transcriptional responses separate antibiotics with different mechanisms of action (MOA’s). **A.** Project setup and overview. Wildtype and adapted strains of *S. pneumoniae* were exposed to different classes of antibiotics, and their fitness outcomes were determined from growth curves. Temporal RNA-Seq data was used to train models that predict the MOA of an antibiotic, and the fitness outcome of a strain. The concept of entropy is developed expanding predictions on fitness to other strains and species and non-antibiotic conditions. CWSI (cell wall synthesis inhibitors): PEN – penicillin, VNC – vancomycin, CEF – cefepime; DSI (DNA synthesis inhibitors): CIP – ciprofloxacin, LVX – levofloxacin; RSI (RNA synthesis inhibitor): RIF – rifampicin; PSI (protein synthesis inhibitors): KAN – kanamycin. TET – tetracycline. **B.** Principal component analysis (PCA) of differential expression datasets (log2FoldChange of +/- antibiotic) from sensitive strains depicts antibiotic responses as largely distinct temporal transcriptional trajectories. **C.** Pairwise distances between PCA trajectories (see Supplementary Methods). Transcriptional trajectories to drugs within the same MOA are similar and tend to cluster together indicated by k-means clustering. **D.** Heatmap of differential expression of an 8-gene panel. Dendrogram shows antibiotic MOA’s can be well separated by differential expression of this gene panel. **E.** Antibiotic MOA prediction by a support vector machine (SVM) trained on differential expression profiles of the 8-gene panel in a training set (Train) and test set (Test). Color intensity is proportional to the number of predictions.

To infer MOA’s from the transcriptional profiles, a regularized logistic regression was used for unbiased feature selection. Since, the number of genes available to use as features far exceeds the number of RNA-Seq experiments this results in a high risk of overfitting, which can be overcome by strong regularization. As previous gene-panel models have used up to 9 genes in any one panel (5, 6), we set the regularization strength such that up to a total of 10 genes would be selected, with at least one gene for each MOA. This reduced the data to a set of 8 genes (2 selected for CWSI, 3 for DSI, 2 for PSI and 1 for RSI) that maximally separate the MOA’s (**Figure 1D**, Supplemental Tables 2, 3). Notably, several genes in this panel and their differential transcription patterns are functionally related to the antibiotics’ MOA. First, up-regulation of the repair DNA polymerase I (SP_0032) is predictive of a DSI which causes double stranded breaks by trapping DNA topoisomerase IV and DNA gyrase. Second, increased transcription of RNA polymerase subunit D (SP_1073) predicts a response to the transcription inhibitor rifampicin (RSI) and may also represent a feedback response to rifampicin to maintain transcription levels. Third, the PSI kanamycin drastically increases transcription of *clpP* (encoding a subunit of Clp protease) by 270-fold and 137-fold in T4 and 19F, respectively. Aminoglycosides cause protein mistranslation, which leads to protein aggregate formation that could be relieved by Clp protease (10). A similar response to PSI was observed in *B. subtilis* (11), indicating that this may be diagnostic of a cellular response to remove mistranslated proteins. Finally, in the presence of CWSI there is up-regulation of the two-component system CiaRH (*ciaR* (SP_0798)), which has been associated with preventing lysis triggered by CWSIs in *S. pneumoniae* (12, 13). In addition to these genes that are clearly connected to a specific MOA, the panel contains genes with less clear connections with the MOA, i.e. SP_2107 (maltose metabolism), SP_2141 (glycosyl hydrolase-like protein), SP_0021 (deoxyuridine 5’-triphosphate nucleotidohydrolase) and SP_2229 (Trp tRNA synthetase) that allow the set to distinguish MOAs. The presence of such genes, along with well-characterized genes relevant to the MOA confirms that the gene panel selection is unbiased and independent of gene function annotations.

To enable MOA predictions from expression data without prior knowledge of the antibiotics used, the 8-gene panel profiles were used to train a support vector machine (SVM) that classifies the MOA of the antibiotic, which resulted in accurate prediction of MOAs (0.92) (**Figure 1E** - Train). The SVM was validated using an independent transcriptional data-set collected from T4 and 19F treated with cefepime (CEF), tetracycline (TET) and ciprofloxacin (CIP; **Figure 1E** - Test, Supplemental Table 1), resulting in an accuracy of 0.75 (random chance results in an accuracy of 0.25). The lower accuracy in the test set (compared to the training set) is mostly due to misclassification of ‘early’ exposure time points or confusion of the model between the PSI and RSI MOA (Supplemental Notes), which can also be seen in the close resemblance between the RSI and PSI PCA trajectories (**Figure 1B**). From a biological perspective, this similarity is likely due to the closely related cellular functions blocked by the two MOAs (i.e. transcription and translation).

### Low and high fitness outcomes after antibiotic exposure or nutrient deprivation of *S. pneumoniae* can be represented by a single 10-gene signature

As T4 and 19F are susceptible to most antibiotics used, the transcriptional profiles in the presence of antibiotics mostly represent cases of low fitness (**Figure 2A**, sensitive strain, 1xMIC_WT_). Besides the MOA-specific responses, we hypothesized that there may also be a single signature of low fitness outcomes that can be extracted from the transcriptional profiles. In order to find patterns that differentiate fitness outcomes (presence or absence of growth in a particular environment), we generated strains with increased fitness in the presence of antibiotics through serial passaging wildtype T4 and 19F in the presence of increasing amounts of antibiotics. Four independent populations for each strain were selected on individual antibiotics to a point that they would grow at 1.5xMIC of the wildtype strain, albeit with a small growth defect (Supplemental Figure 1A, B). To identify signatures of fitness, the wild type and adapted clones were exposed for 2 to 4 hours to 1x and 1.5-2xMIC of the antibiotics used for adaptation, followed by RNA-Seq at different time points after exposure (**Figure 2A**, Supplemental Figure 1A). In parallel, RNA-Seq was performed on *S. pneumoniae* strains D39 and T4 in a chemically defined medium, and media from which either uracil, Glycine or L-Valine was removed. This allowed for the identification of a common stress signature that applies to nutrient deprivation as well, as these nutrients are essential for D39 but not T4. Lastly, D39 was adapted to grow in the absence of each individual nutrient, after which RNA-Seq was repeated for adapted clones (Supplemental Table 1 lists all 24 strains, 67 populations and 6 species, and 267 RNA-Seq experiments; visualizable and explorable online at http://bioinformatics.bc.edu/shiny/ABX).

**Figure 2.**
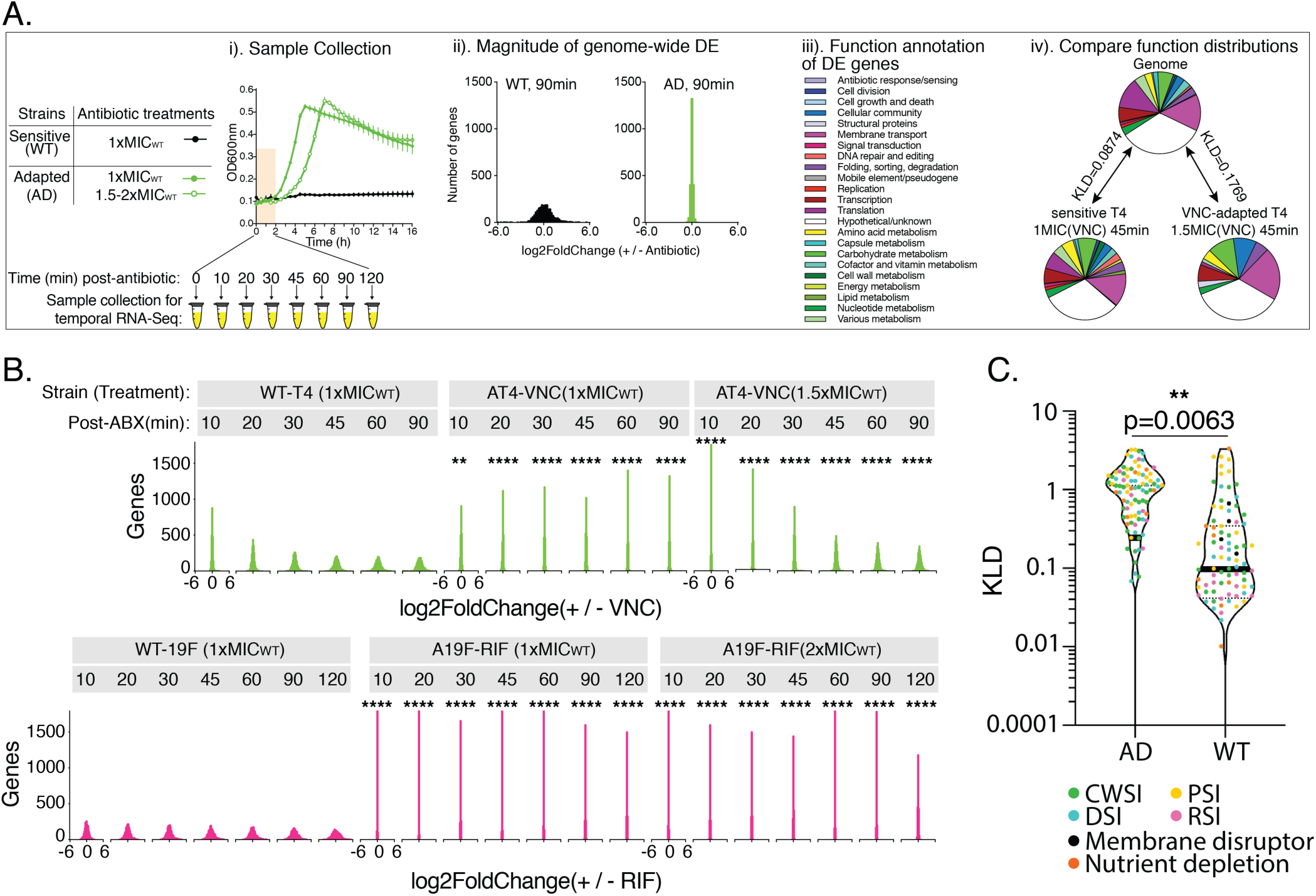
A transcriptional response separates bacteria with different sensitivities to the same antibiotic. **A.** A schematic illustration of temporal RNA-Seq sample collection (i) and data processing (ii-iv) on stress-sensitive wild-type (WT) and stress–insensitive adapted strains (AD), which are obtained through experimental evolution of the WT strains. Magnitude distribution of genome-wide differential expression is compared between each pair of WT and AD strains (ii). Genes in the *S. pneumoniae* genomes are divided over 23 gene function categories (see figure legend). Kullback-Leibler divergence (KLD) of gene function distributions between genes with significant expression changes and function distribution of all genes present in the genome (iii). A similar function distribution to the genome is indicated by a low KLD value, e.g. T4-VNC(1xMIC)-45min, while a dissimilar function distribution is indicated by a high KLD value, e.g. VNC-adapted T4-VNC(1.5xMIC)-45min (iv). **B.** The magnitude of genome-wide differential expression (indicated as log2FoldChange Antibiotic/NDC (no drug control)) shows significantly wider distributions in antibiotic-sensitive strains (wtTIGR4 and wt19F) compared to antibiotic-adapted strains in the presence of vancomycin (a cell wall synthesis inhibitor; CWSI) and rifampicin (an RNA synthesis inhibitor; RSI), respectively in a Kolmogorov-Smirnov test. *: 0.001<p<0.05; **: 0.0001<p<0.001; ***: p<0.0001. See **Supplemental Figure 2D, E** for other antibiotics. All histograms are on the same scale of −6 to **C.** AD have a significantly higher KLD than WT in the presence of antibiotics or absence of D39-essential nutrients in an unpaired t-test.

To determine whether specific transcriptional differences exist that distinguish a strain that successfully grows in a particular environment (high fitness) from one that does not (low fitness), the dataset was split into stress-sensitive and stress-insensitive (e.g. adapted) groups. This approach identified three transcriptional patterns. First, an unadapted, stress-sensitive strain tends to trigger a greater number of transcriptional changes at 1xMIC or during nutrient deprivation compared to an adapted strain (Supplemental Figure 2A-C). Second, in response to antibiotic or nutrient deprivation stress, the magnitude of transcriptional changes in the stress-sensitive strains shows a broad distribution over time compared to the adapted strains, indicating that there is extensive genome-wide transcriptional change in sensitive strains (**Figure 2A, B**; Supplemental Figure 2D-F). Finally, stress-sensitive strains have a significantly lower Kullback-Leibler divergence (KLD) relative to adapted strains, indicating higher similarity of differentially expressed (DE) genes compared to the distribution of the entire genome. A lower KLD therefore indicates a similar function distribution in DE genes and the genome, which can be interpreted as a lack of functional enrichment in the differentially expressed genes (**Figure 2A, C**).

Logistic regression was used on the training part of the RNA-Seq dataset (231 experimental conditions; Supplemental Table 1) to identify a gene panel that could accurately separate stress-sensitive from stress-insensitive responses in a strain- and environment-independent manner. Similar to the selection of the MOA-prediction gene panel, regularization was applied such that the number of genes used did not exceed 10. This resulted in a 10-gene-panel (Supplemental Tables 2, 3) that is able to separate and predict low and high fitness outcomes with an accuracy of 0.91 for the training set (**Figure 3A**) and 0.83 for the independent test set (**Figure 3B**, Supplemental Table 4; see Supplemental Notes for details on misclassifications). These 10 genes showed similar DE patterns in low fitness outcomes in all the tested strains and environments, and remained mostly unchanged in high fitness outcomes (**Figure 3C**). The functions of these genes include cell division (*ftsZ*; SP_1666), metal ion transport (SP_1857 and SP_1869), competence (SP_0955), cell surface protein (SP_2201), transcriptional regulation (SP_1856 and SP_1800), translation (SP_0929) and metabolism (SP_1478 and SP_0589). Unlike the MOA gene panel, genes in the fitness gene panel are not obviously related to the antibiotics or depleted nutrients, but may be pointing to a more general stress-response.

**Figure 3.**
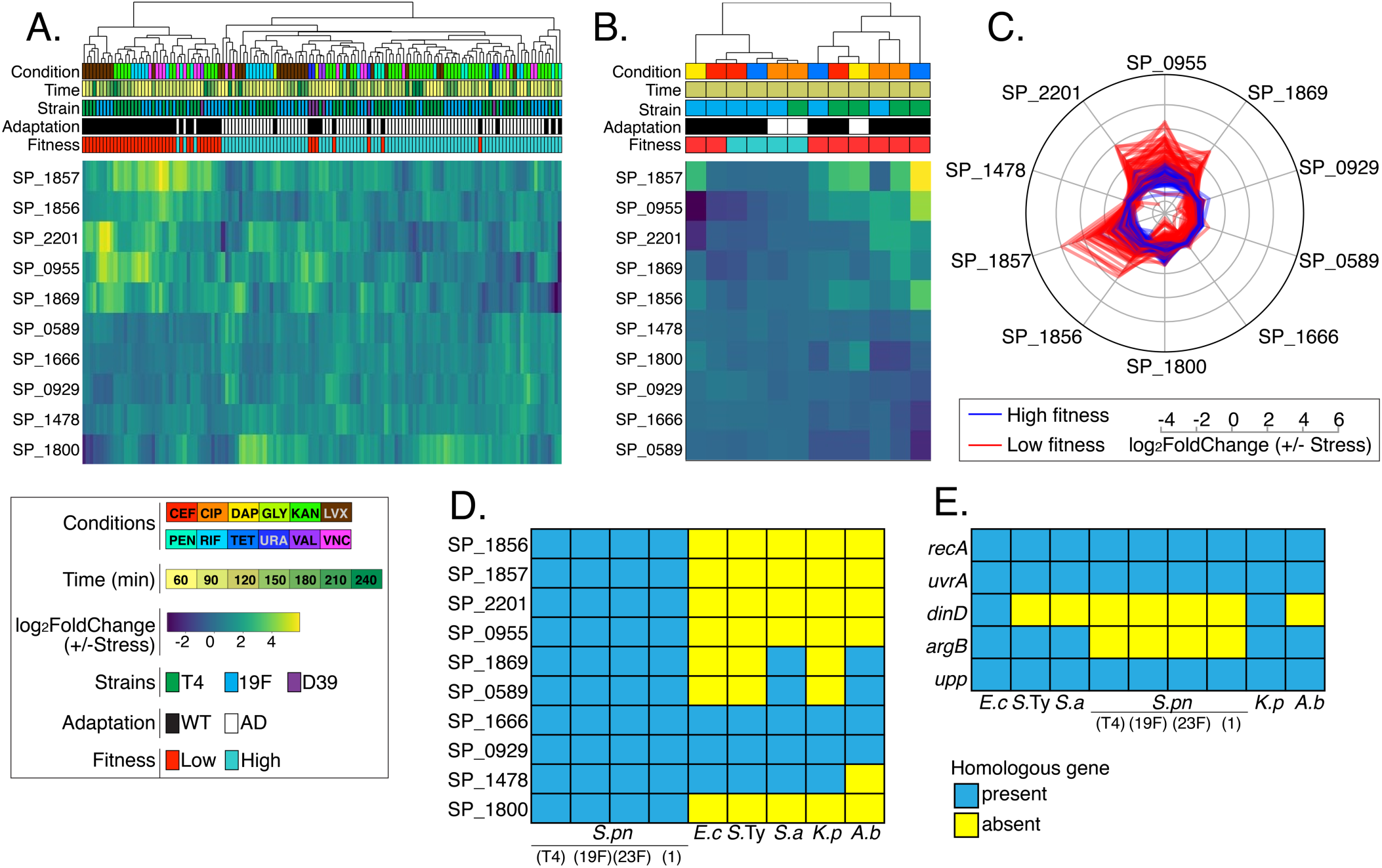
A 10-gene panel predicts fitness outcomes of *S. pneumoniae* under antibiotic and nutrient stress. A 10-gene panel is generated by logistic regression on 231 RNA-Seq profiles collected from antibiotic exposure or D39-essential nutrient depletion conditions in stress-sensitive and stress-insensitive strains. Log2fold change (+/- stress) of the 10-gene panel for fitness is indicated by the heat-map color gradients, which separates high fitness (blue in the ‘Fitness’ bar) from low fitness (red) in a training (**A.**) and test set (**B.**). **C.** Differential expression of each gene in the gene panel is depicted as a radar plot, showing a clear difference between low (red) and high (blue) fitness outcomes. Presence and absence of each gene in the *S. pneumoniae* fitness panel (**D**.) or *E. coli* ciprofloxacin sensitivity panel (**E**; (5)) in six pathogenic species is first determined by protein BLAST based on three criteria: query coverage > 50%, E value < 1E-50 and percent identity > 30%, indicating that genes in a gene panel are poorly conserved and thus might suffer a limited generalizability across multiple species. *S.pn*: *S. pneumoniae, E.c*: *E. coli, S.*Ty: *S.* Typhimurium, *S.a*: *S. aureus, K.p*: *K. pneumoniae, A.b*: *A. baumannii*

### Transcriptional responses can be captured by entropy, a generalizable feature for fitness predictions

Although the gene panel-based fitness predictions show high accuracy for the tested conditions in *S. pneumoniae*, it is unlikely that a small gene panel model is able to predict fitness outcomes for a wider variety of conditions or for multiple species. Indeed the 10 *S. pneumoniae* genes that indicate fitness outcomes are poorly conserved in *E. coli, Salmonella* Typhimurium and 3 ESKAPE species, including *Acinetobacter baumannii, Klebsiella pneumoniae*, and *Staphylococcus aureus* (**Figure 3D**). Similarly, only 3 out of 5 genes in a published *E. coli* ciprofloxacin sensitivity gene panel are present in all 6 species, indicating that a gene panel approach is limited in its generalizability ((5); **Figure 3E)**.

Our analysis indicates that in low-fitness cases, there is a corresponding disruption in transcriptional regulation compared to a high-fitness scenario. To explore this idea, we leveraged the concept of entropy to quantify the extent of transcriptomic disruption triggered by a stress. Using the information theory definition of entropy, this concept captures the amount of chaos in a system, implemented here by quantifying the variance in differential expression that occurs over time under stress conditions in a gene or a collection of genes. (**Figure 2B**, Supplemental Figure 2). Our dataset contains 29 detailed RNA-Seq time course experiments for which entropy was calculated initially by taking each gene in the system as an independent entity (**Figure 4A**, Model 1). The average of the extent of transcriptional perturbation across all genes in the genome was quantified by defining entropy as the average of the variance in differential expression over time for all genes (Supplemental Methods, Equation 2; **Figure 4A**, Model 1). To enable predictions, a model based on this single feature was trained by finding just one scalar number, a threshold for entropy, to distinguish high fitness (below threshold) from low fitness (above threshold) (**Figure 4B**, Supplemental Figure 3B). This model performs well, with only 1 false negative, i.e. a sample with high fitness having an entropy value above the threshold (**Figure 4B**), but it assumes that each gene behaves independently. This assumption does not capture biological complexity, as many genes co-vary in expression, which can lead to over-estimating entropy if these dependencies are not accounted for.

**Figure 4.**
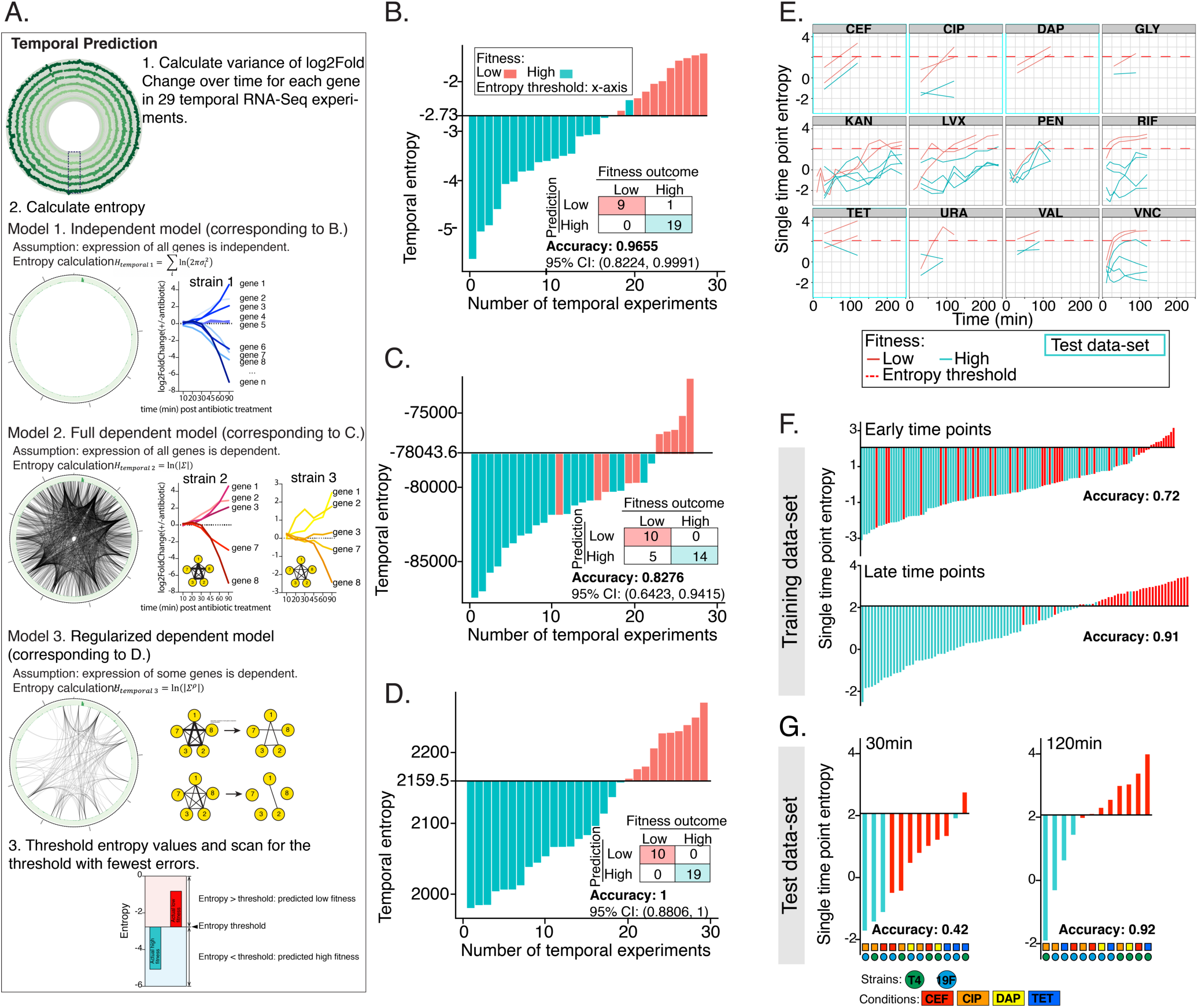
Temporal and single time point entropy calculated from a transcriptional response predicts high and low fitness outcomes. **A.** Illustration of the three temporal entropy models (Models 1-3) that are applied on 29 temporal RNA-Seq experiments. **B.-D.** Temporal fitness prediction is shown as a ranked plot of entropy in each temporal RNA-Seq dataset. The y-intercept indicates the entropy threshold, i.e. entropy higher than threshold is predicted as a low fitness outcome; entropy lower than threshold is predicted as a high fitness outcome. A confusion matrix is generated for each model by comparing to the actual fitness outcome (High/low fitness: cyan/red bars). **E.** Single time point entropy is calculated from differential expression of all genes in an experiment at one time point and plotted against time post-stress exposure (presence of antibiotics – CEF, CIP, DAP, KAN, LVX, PEN, RIF, TET, VNC, or absence of nutrients – Glycine-GLY, Uracil-URA, Valine-VAL). Dashed red line indicates the entropy threshold for this model. Training and test sets are indicated by grey or blue borders respectively. Fitness prediction is performed at each single time point (231 data points) in the 29 temporal experiments as the training set (**F.**), and validated by an independent set of single time point RNA-Seq experiments as the test set (24 data points; **G.**). In both training (**F.**) and test (**G.**) datasets, fitness prediction is more accurate from late time points than early time points indicating late time points are more characteristic of a strain’s fitness outcome.

To correct for co-variances in expression and their potentially confounding effects, we consider a gene co-expression network, where, if the transcription of two genes have high co-variance, these genes are connected. While the underlying co-transcriptional network is unknown for *S. pneumoniae*, it can be inferred using the available temporal RNA-Seq data by computing the inverse of the co-variance matrix of transcriptional changes among all gene pairs. This approach yielded a complete network, in which all possible links between gene pairs were present and weighted by the co-variance values. This is equivalent to considering the differential expression of all genes from all time points to be a multivariate Gaussian distribution with as many dimensions as there are genes (Supplemental Methods; Equation 3). We computed entropy for this multivariate distribution and set the appropriate threshold for separating high and low fitness (**Figure 4A**, Model 2). All samples with higher entropy than the threshold were instances of low fitness (**Figure 4C**, Supplemental Figure 3C), but 5 low fitness samples appeared to have low entropy as well (false positives). This result is likely a consequence of considering ‘raw’ co-variance values, which leads to the appearance of spurious links in the network that do not reflect real links between genes. Therefore, Model 2 may overestimate the number of links between genes. To correct for such spurious links, regularization was applied on the inverse of the co-variance to obtain a sparse co-transcriptional network. Note that this model (**Figure 4A**, Model 3) is equivalent to Model 1 if regularization is very stringent (all links between genes are dropped), and equivalent to Model 2 if no regularization is applied (all links are present). By adjusting the level of regularization, we show that temporal entropy can reach an accuracy of 1 (**Figure 4D**, Supplemental Figure 3D, E, Supplemental Table 5), demonstrating the utility of this single feature in classifying fitness outcomes.

The time course experiments accurately capture a bacterium’s survival in a test environment, but they are labor intensive and potentially expensive. Therefore, a single time point prediction model was trained, by quantifying the variance of the DE magnitude distribution for all single time points as a measure of entropy (Supplementary Methods, Equation 1). Similar to the temporal models, the only parameter that we fit was the threshold for entropy (in this case 2.07), which is the value that maximizes classification accuracy in the training set. Analogous to the temporal models, low fitness is associated with higher entropy than high fitness conditions (**Figure 4E**, Supplemental File 2). Importantly, cases of high and low fitness are better separated at later time points (accuracy = 0.91 in the training and 0.92 in the test dataset) than at early time points (accuracy = 0.74 in the training and 0.42 in the test dataset), indicating that time points from the second half of the time course experiments are more characteristic of fitness outcome than those from the first half (**Figure 4F**). Specifically, when applied on an independent test dataset, the single time point entropy model led to 7 false positive predictions based on the early time points even though the misclassified low fitness cases have a higher overall entropy than that of high fitness cases (**Figure 4G**). This suggests that it is possible to successfully train an early time point-specific model if more training data were used (Supplemental Figure 3A). In contrast, the single time point entropy predictor, with the same threshold value of 2.07, performs very well for the late time points on the test dataset, misclassifying only 1, that was very close to the threshold, out of 12 experiments (**Figure 4G**).

### Entropy-based fitness predictions are strain, species and stress-type independent and can be used to infer the antibiotic minimum inhibitory concentration (MIC)

To test if the entropy-based approach is generalizable and extends to other *S. pneumoniae* strains or other species, a new RNA-Seq dataset was generated to predict fitness outcomes under ciprofloxacin exposure for *Salmonella* Typhimurium, *S. aureus, E. coli, K. pneumoniae* and two additional *S. pneumoniae* strains representing serotypes 1 and 23F (Supplemental Table 1). These five species represent both Gram-negative and Gram-positive bacteria and cover a wide range of ciprofloxacin MICs (**Figure 5A**). The overall response characteristics are similar to what was observed for *S. pneumoniae*, with 120 minutes exposure to 1µg/mL ciprofloxacin triggering an expansion of expression changes from bacterial cultures having low fitness (*S*. Typhimurium and *S. pneumoniae* serotype 1), compared to those with high fitness (*S. pneumoniae* serotype 23F, *E. coli* and *K. pneumoniae*) (**Figure 5B**). When exposed to a higher dose of CIP (strain-specific 1xMIC_CIP_) the organisms with lower CIP-sensitivity were shown to trigger an increased number of expression changes with a wider magnitude, indicative of their lowered fitness at the increased concentration (**Figure 5B**, *S. pneumoniae* serotype 23F, *E. coli* and *K. pneumoniae*). Next, each transcriptional response was captured by an entropy calculation. Importantly, with the original threshold of 2.07 we had calibrated with data from *S. pneumoniae* in Figure 4, fitness outcomes could be predicted for the new organisms with 100% accuracy, indicating that the amount of transcriptional disruption by antibiotics is a species-independent generalizable feature for fitness outcome. Therefore, entropy could be visualized graphically (**Figure 5B**), and quantified to make high-accuracy predictions.

**Figure 5.**
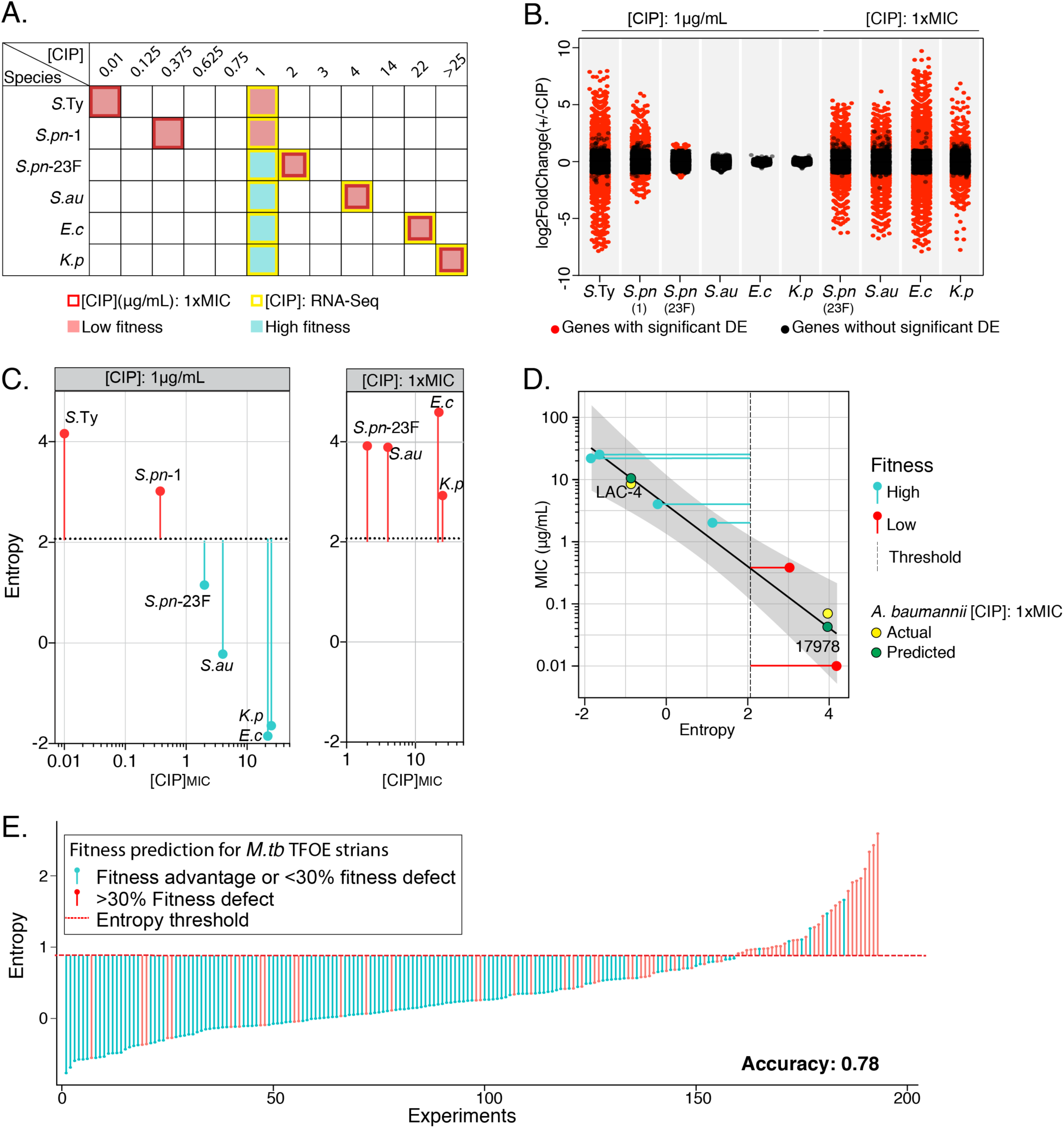
Entropy based fitness predictions extend to multiple species under antibiotic and non-antibiotic stress. **A.** Six strains representing 5 species are ranked from low to high ciprofloxacin minimal inhibitory concentrations (MIC_CIP_) tested by growth curve assays. The multi-species CIP RNA-Seq is performed at two CIP concentrations: 1) 1µg/mL for all 6 strains corresponding to 2 low fitness outcomes (red squares) and 4 high fitness outcomes (cyan squares); 2) MIC_CIP_ for strains that are insensitive to 1µg/mL of CIP, i.e. *S. pneumoniae* serotype 23F, *S. aureus* UCSD Mn6, *E. coli* AR538, and *K. pneumoniae* AR497, corresponding to 4 additional low fitness outcomes. The number of genes that change in expression upon exposure to 1µg/mL CIP (|log2FoldChange|>1 and p-adj<0.05) as well as their change in magnitude is inversely correlated to their CIP sensitivity (**B.**) and their entropy (**C.**). Additionally, strains with MIC_CIP_ higher than 1µg/mL revert to triggering a large number of differential expression genes (**B.**) and a high entropy (**C.**) at their respective 1xMIC_CIP_. **D.** Using a linear regression model (black line; error band: 95% CI), MIC’s are predicted for *A. baumannii* strains ATCC 17978 and LAC-4 based on their entropy at 1µg/mL of ciprofloxacin. The predicted (black) and measured (red) MIC for the two strains are similar for both strains. See **Supplemental Figure 1D** for MIC determination for *A. baumannii* ATCC17978 and LAC-4. **E.** Entropy calculated from transcriptional profiles of 193 *M. tuberculosis* transcription factor over-expression (TFOE) strains separates strains with a >30% fitness defect upon TFOE induction (red) from strains with a fitness advantage or <30% fitness defect upon induction (cyan). At the threshold of 0.71 (red dotted line), fitness outcomes are correctly predicted at an accuracy of 0.78.

Interestingly, the entropy measurement of each strain was found to be inversely proportional to the MIC_CIP_ (**Figure 5C**), consistent with transcriptional disruption being proportional to stress sensitivity. The correlation between entropy and ciprofloxacin sensitivity in **Figure 5C** (left panel) therefore implies that the antibiotic sensitivity of any species could be predicted from its transcriptomic entropy. To test this, entropy was calculated for *Acinetobacter baumannii* isolates that are either low (ATCC 17978) or high (LAC-4) virulence, by collecting RNA-Seq profiles after 120 min exposure to 1μg/mL of ciprofloxacin. Using a linear regression model, the ciprofloxacin MICs of the *A. baumannii* strains were predicted to be 0.04 and 10.45μg/mL, which are proximate to the measured MIC’s of 0.07 and 8.5μg/mL for ATCC 17978 and LAC-4, respectively (**Figure 5D**; Supplemental Figure 1D). This demonstrates that entropy can be applied to determine antibiotic sensitivity for new species, and is not simply a binary indicator of fitness outcomes.

To explore the applicability of entropy beyond nutrient and antibiotic stress, we performed entropy-based fitness classification on a published collection of 193 *M. tuberculosis* transcription factor over-expression (TFOE) strains (14). Upon TFOE, these strains exhibit fitness changes, ranging from severe growth defects to small growth advantages (15). Over-expression of a single transcription factor can thereby exert stress on the bacterium that can result in drastically different fitness outcomes. By calculating entropy from whole-genome microarray data collected from each TFOE strain, using genome-wide DE under inducing and noninducing conditions, it is possible to distinguish strains based on their fitness levels at an accuracy of 0.78, using a newly trained entropy threshold for this dataset (**Figure 5E**). This result compares favorably with a more complicated approach involving the integration of each TFOE transcriptional profile into condition-specific metabolic models (14). These data show that entropy has the potential to be utilized as a strong and generalizable fitness prediction method for both antibiotic and non-antibiotic stress, using different data types and a large variety of bacterial strains and species.

### A systems-level view successfully predicts fitness and is less sensitive to noise from a single feature

While the transcriptome-based entropy-approach is a strong predictor of fitness, it is a relatively coarse method as it captures only a single organizational level of the response; the transcriptome. In most biological systems, changes occur across multiple levels, which are detected by alternative readouts, such as identification of phenotypic changes resulting from mutations (16-21). Furthermore, entropy ignores details present in the data, such as the type of stress and the functional roles of genes, that may increase the accuracy or widen the applicability of the approach. For instance, within the transcriptomic data, different gene sets show different levels of perturbation, as witnessed in *S. pneumoniae* T4, in which genes in the category “Antibiotic Response and Sensing” show higher entropy than other functional categories (**Figure 6A**, Supplemental Figure 4A). In contrast, essential genes are mainly down-regulated and nonessential genes are evenly distributed as up- and down-regulated in the antibiotic challenge conditions (**Figure 6B**, Supplemental Figure 4B).

**Figure 6.**
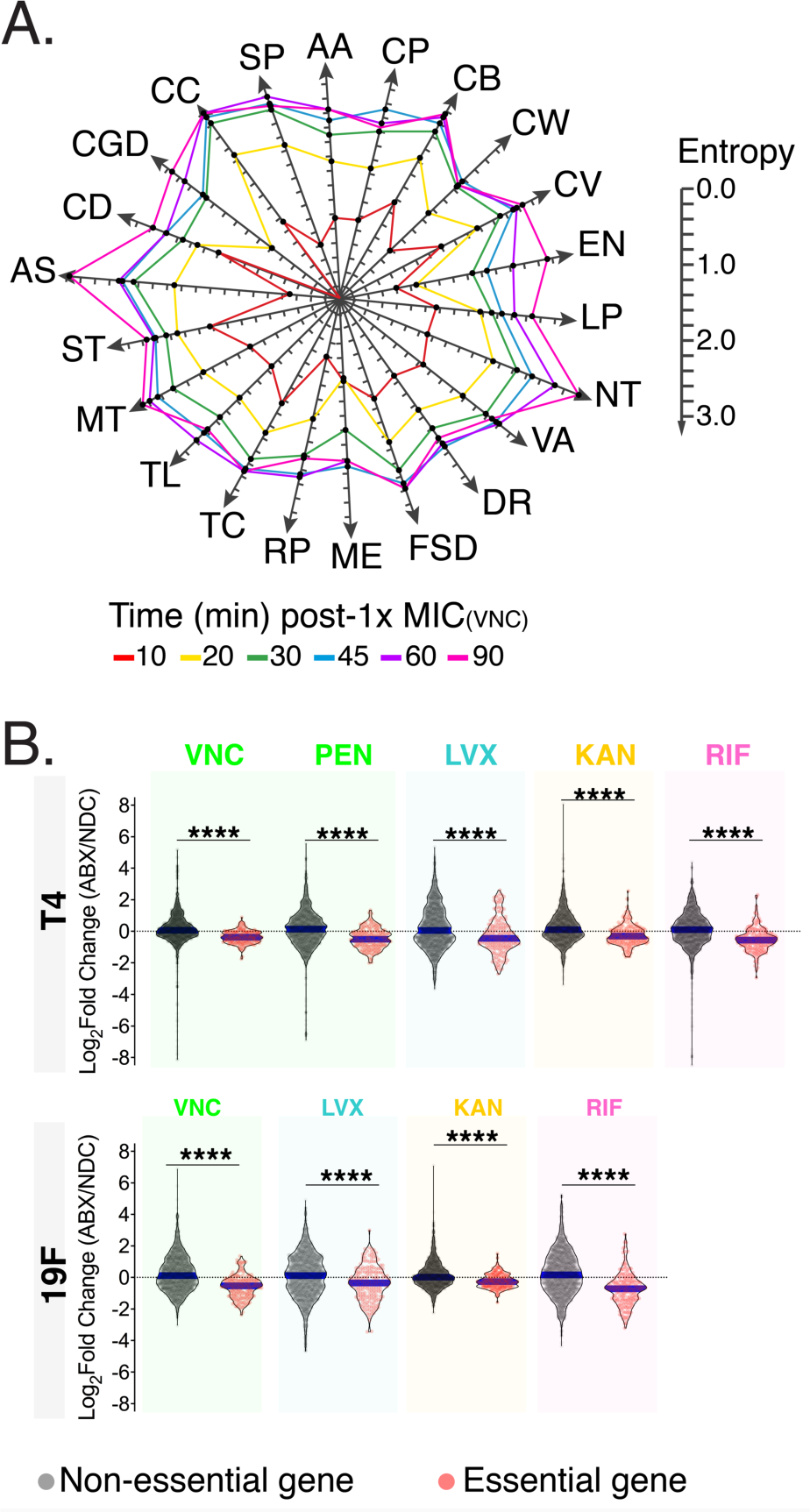
Detailed features can be extracted from transcriptomic responses of strains with low and high fitness outcomes. **A.** The entropy of a transcriptional response can be split by gene function to indicate the level of transcriptional perturbation in each cellular system. AA: amino acid metabolism, CP: capsule metabolism, CB: carbohydrate metabolism, CW: cell wall metabolism, CV: cofactors and vitamin metabolism, EN: energy metabolism, LP: lipid metabolism, NT: nucleotide metabolism, VA: various metabolism, DR: DNA repair, FSD: folding, sorting and degradation, ME: mobile elements. RP: replication, TC: transcription, TL: translation, MT: membrane transport, ST: signal transduction, AS: antibiotic sensing, CD: cell division, CGD: cell growth and death, CC: cellular community, SP: structural proteins. **B.** Essential genes tend to be significantly more down-regulated compared to non-essential genes in stress-sensitive strains (wt-T4; unpaired t-test, *: 0.001<p<0.05; **: 0.0001<p<0.001; ***: p<0.0001).

In order to make use of these detailed observations, we assembled a set of 54 features that describe various aspects of a response, the organism and the experienced stress. To this end, we considered the type of stress, such as the MOA of an antibiotic, the phenotypic response to stress as characterized by Tn-Seq, and the transcriptional response (**Figure 7A**; Supplemental File 3). This approach yields a large number of features relative to the number of samples (54 features and 231 samples in the training set), so we performed a round of feature selection using a regularized logistic regression model (**Figure 7A**). The six most informative features (described in Supplemental Methods) were used to train an SVM that is able to classify the fitness outcomes with high accuracy, by setting a threshold in which a probability >0.5 translates to high fitness, and probability <0.5 to low fitness (**Figure 7B**, Supplemental File 2, Supplemental Figure 4C). Importantly, accurate predictions could be made at both early and late time points for antibiotics that triggered a fast transcriptomic response, as low and high fitness outcomes were well-separated as early as 20 min after rifampicin exposure and 30-45 min after vancomycin exposure (**Figure 7B**). For other antibiotics, accurate predictions could only be made at later time points. For example, kanamycin treatment resulted in false positives up to and including 120 min, possibly due to a slow transcriptional response to this drug (**Figure 7B**; Supplemental Figure 2A, D, E -Kanamycin). The primary advantage of this model is that it provides less ambiguous predictions compared to the predictive models based solely on transcriptional entropy, with the prediction probability following a bimodal distribution that has very few cases near the threshold (= 0.5) in the training or independently generated test sets (**Figure 7C**). Furthermore, in the test set all misclassified cases are at the early (30 min) time point, while 100% of the late time points (120 min) are classified correctly (**Figure 7D**). This classifier (test set accuracy = 0.5 and 1 in early and late time points respectively) outperformed the single-time point entropy classifier (test set accuracy = 0.42 and 0.92 in early and late time points respectively). Thus, by increasing data resolution and including multiple data sources, a highly accurate model is achieved that is robust to noise from a single data source.

**Figure 7.**
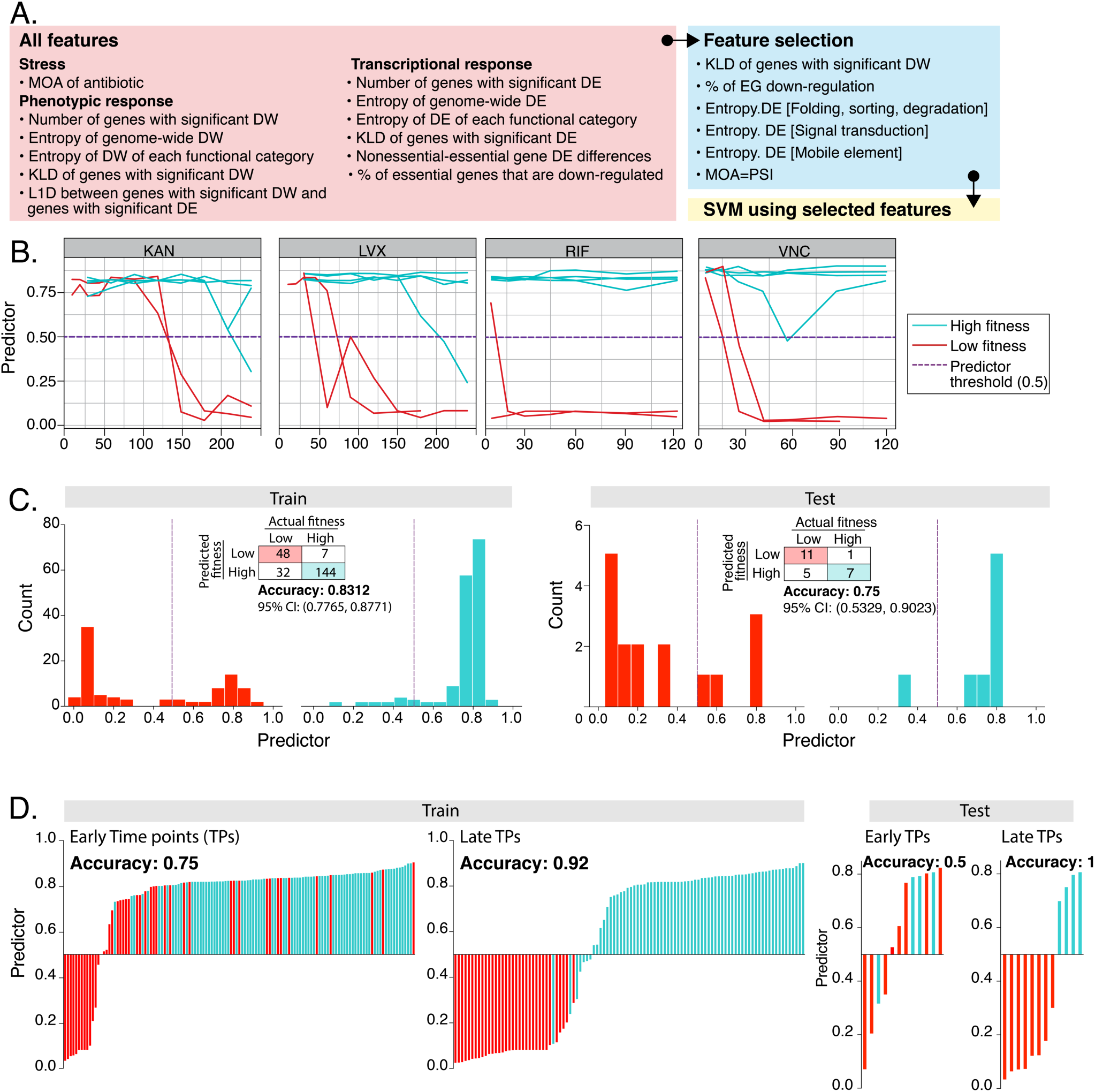
A systems-level view of the bacterium further improves fitness predictions. **A.** Illustration of the construction of a complex feature classifier (CFC) in three main steps: 1) assembly of all input features, 2) feature selection and 3) SVM-based fitness predictions. **B.** Prediction probabilities generated by the CFC are plotted at each time point for strains with high fitness (cyan line) and low fitness (red line) in the presence of KAN, LVX, RIF, and VNC. A probability higher than the threshold (0.50; red dotted line) is predicted as high fitness; while a predictor lower than the threshold is predicted as low fitness. **C.** Performance of the CFC on training and test datasets is shown as frequency distributions of prediction probabilities separated by actual fitness outcomes and the corresponding confusion matrices. **D.** High (cyan) and low (red) fitness outcomes are mostly well separated by the CFC predictor at the threshold of 0.50. Similar to the single time point entropy model (Figure 3), fitness prediction by the CFC achieves a much higher accuracy of 1.0 at late time points compared to early time points.

## Discussion

A major goal of this work was to determine if there is a quantifiable feature in the bacterial environmental response that can accurately predict fitness in that environment that is independent of strain, species or the type of stress. To be generalizable, the selected feature needs to be common across species and environments. By generating a large experimental dataset we discovered that such a feature exists, namely transcriptomic entropy, which represents the level of transcriptional disruption that occurs in a system while responding to the environment. Centering on entropy, we develop a suite of statistical models that vary in their complexity and that accommodate different types and amounts of input data enabling predictions on the MOA of a stress, the fitness outcome of bacteria in a variety of different environments, and the MIC.

The developed fitness prediction models differ in approach and input data required. The first model extracts a small set of informative features from genome-wide transcriptional profiling data (i.e. a gene panel). Although we have shown previously that phenotypic change and expression change rarely overlap on the same genes, there are cases in which a gene can be simultaneously transcriptionally and phenotypically important under a stressful condition (16). Indeed, several members of the fitness gene panel are required for bacterial growth. For example, inhibition of the essential cell division gene *ftsZ* has been shown to cause growth inhibition of MRSA (22). Among the non-essential genes, transposon insertion mutants in SP_0929, SP_0589 and SP_1856 resulted in a significant fitness decrease in the presence of multiple antibiotics (LVX, TET, CIP, CEF and RIF; Supplemental File 1). Therefore, further characterization of the genes in this panel might reveal potential antimicrobial targets. Second, by applying a first-principle approach, a single feature (entropy) is defined to capture an intuitive property of a transcriptome: the extent of perturbation. Since entropy does not rely on responses in specific genes, entropy-based models extend beyond a single species. Moreover, the single timepoint entropy can be used in a regression model, offering a finer resolution by predicting the level of sensitivity to a particular stress (in this case ciprofloxacin) as well. Finally, a third approach is a data-driven one, in which genome-wide data across multiple biological systems were utilized (i.e. SVM for fitness prediction). Importantly, all three types of fitness prediction models underscore that a bacterium’s fitness phenotype is predictable, and accurate predictions can be made in multiple pathogens independent of the mechanisms of stress, including different classes of antibiotics and nutrient depletion.

By demonstrating the feasibility of predictions of fitness outcomes and antibiotic sensitivity, we believe this work could provide novel opportunities to contribute to infectious disease diagnostics such as antibiotic susceptibility testing (AST). Although AST can be completed in a relatively short amount of time for many pathogenic species, it remains a lengthy process for slow-growing species such as *M. tuberculosis* (23). Therefore, it is desirable to be able to predict the fitness outcome of such slow-growing species as early as possible, for instance using RNA expression data. RNA-based detection has previously been applied to correlate antibiotic susceptibility with the expression of several genes (5, 6, 24). However, due to variability in gene-homology, such approaches can rapidly become species specific, and even when a gene is present, the way it responds to a stress might not be the same. For instance, among the nine strains and species we sampled here, the presence/absence of gene’s in the 10-gene panel for fitness predictions and their expression profiles are highly variable (Supplemental Figure 5A). Additionally, a published *E. coli* ciprofloxacin sensitivity panel is highly specific for that species due to variability in presence/absence and expression patterns (Supplemental Figure 5B; (5)). This indicates, that gene-panel approaches may indeed quickly become strain, species and stress-type dependent. In contrast, we show that entropy-based fitness predictions are independent of gene homology, which allows for generalizability across different species under various stress-types (i.e. antibiotics, nutrient depletion and transcription induction). Second, rather than being a binary prediction method, entropy can be applied to predict the level of antibiotic sensitivity in a species-independent manner, which is useful to determine a bacterium’s level of susceptibility to an antibiotic without performing possibly more extensive growth-based assays or identifying the resistance genes or mutations by whole-genome sequencing. Importantly, we show that MIC predictions can be achieved by profiling the transcriptome of a bacterium at a single time-point and a single concentration of an antibiotic, requiring little prior knowledge of antibiotic MOA or direct targets in the tested species. Our entropy-based antibiotic sensitivity prediction could therefore contribute to improving the speed and generalizability of existing AST methods.

Since transcriptional entropy captures information at a single level, several improvements could be implemented to potentially enhance the fitness prediction model. We show that some antibiotics trigger a faster response while some trigger a slower response (Supplemental Figure 2A, D, E). Consequently, it would make more sense to apply a fitness-predictor to the transcriptomic profile at a later timepoint for antibiotics that trigger a slower response such as kanamycin; but an earlier timepoint can be used for antibiotics such as rifampicin. Knowing (or predicting) the type of antibiotic being used can therefore inform when to use a fitness predictor, i.e. fitness predictions can potentially be enhanced when the MOA and fitness predictors are used in tandem. As indicated by the data-driven complex feature model, fitness predictions are improved when fine-grained information on the bacterial stress response is included. Additional types of data as well as the consideration of more features may thus improve fitness predictions through the inclusion of more detailed information. Thus, by gathering and integrating information pertaining to the host environment, for instance by simultaneous transcriptomic profiling via dual RNA-Seq(25) and cytokine profiling of the host response, our model might be able to infer and monitor disease progression *in vivo*.

To conclude, with entropy we present a novel concept that is independent of gene-identity, gene-function, and type of stress, and can be applied as a fundamental building block for generalizable statistical models that accurately predict bacterial fitness and MICs for Gram-positive and negative species alike.

