## Supplemental Information for "Entropy of a bacterial stress response is a generalizable predictor for fitness and antibiotic sensitivity"

† Equal contribution
* Corresponding author

**This PDF file includes:**

Supplemental Figures 1-5

Supplemental Tables 1-6

Supplemental Notes

Supplemental Methods

Supplemental References

**Other supplemental datasets for this manuscript include the following:**

Supplemental File 1

Supplemental File 2

Supplemental File 3

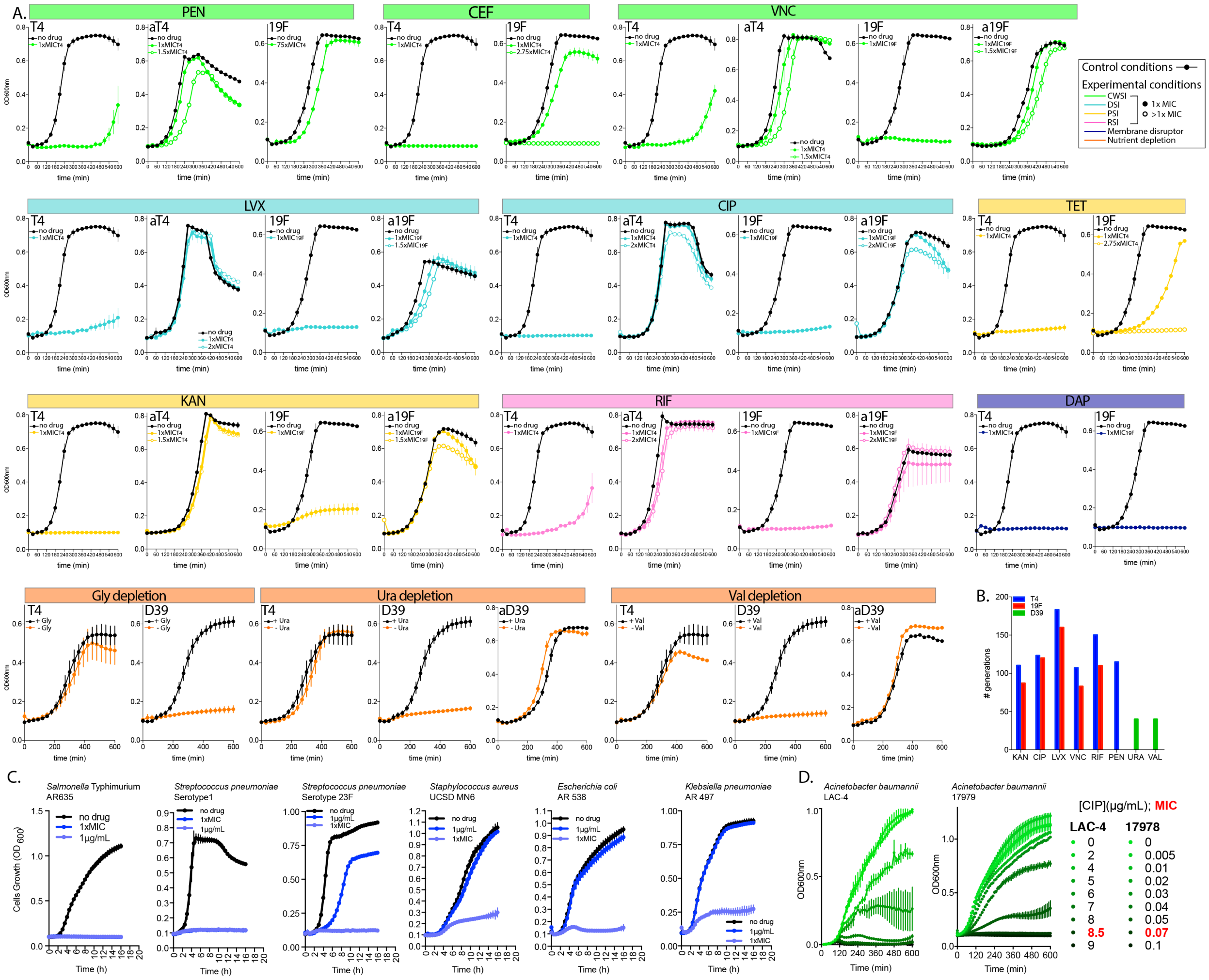


**Supplemental Figure 1.** High and low fitness outcomes under antibiotic exposure and single nutrient depletion.

**A.** Growth curves of stress-sensitive *S. pneumoniae* T4 and 19F strains and antibiotic- or nutrient-adapted strains (labeled as aT4, a19F and aD39). Error bars: standard error of at least three biological replicates.

**B.** Number of generations of adapted populations. Detailed information on minimum inhibitory concentrations is listed in **Supplemental Table 8**.

**C**. Growth curves of *S*. Typhimurium, *S. pneumoniae* serotypes 1 and 23F strains, *S. aureus*, *E. coli* and *K. pneumoniae* under 1μg/mL and strain-specific minimum inhibitory concentration (1xMIC) of ciprofloxacin.

**D.** Growth curves of ciprofloxacin MIC determination for *A.baummannii* strains LAC-4 and ATCC 17978.

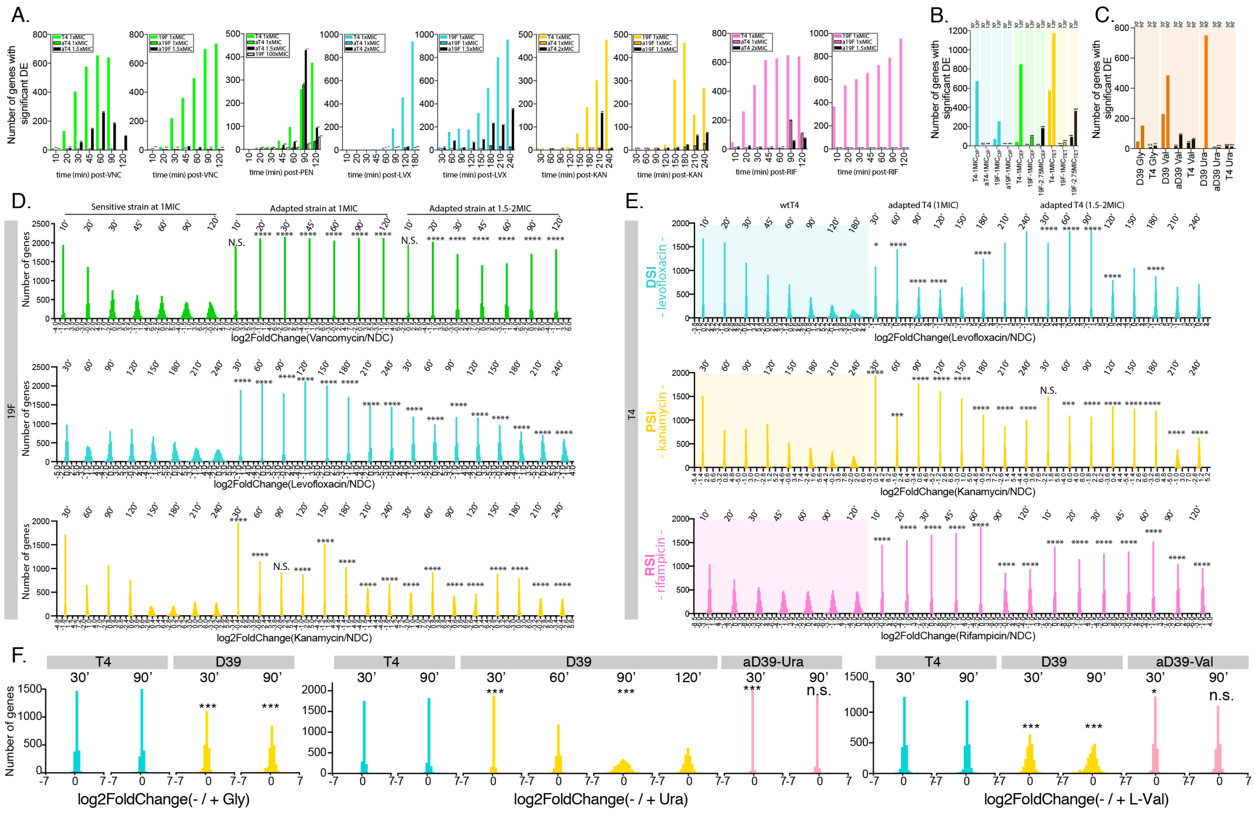


**Supplemental Figure 2.** Transcriptomic signatures distinguish a strain that successfully grows in its environment (high fitness) from one that does not (low fitness).

Numbers of genes with significant differential expression (DE) are mostly significantly higher in sensitive T4 or 19F than in antibiotic-adapted T4 or 19F in the temporal RNA-Seq experiments of VNC, LVX, KAN, RIF, and PEN exposure (**A**) and in the single time-points RNA-Seq experiments of CIP, CEF and TET exposure (**B**). 19F is used as an insensitive strain in CEF and TET experiments, as its growth is not significantly affected by 1x MIC$_{T4}$ of CEF and TET (as shown in **Supplemental Figure 1A**).
**C.** Under single depletion of three D39-essential nutrients (Gly – Glycine, Val – L-Valine, Ura – uracil) the nutrient-dependent D39 triggers significantly more differential expression than the independent T4 or the Val- or Ura-adapted D39. For **A-C**, ****: p<0.00001, ***: 0.00001<p<0.0001, **: 0.0001<p<0.001, *: 0.001<p<0.05 in a two proportion Z-test. Significant differential expression is defined by two criteria: |log2FoldChange(+/-stress)|>1 and padj<0.05.
The magnitude of genome-wide differential expression shwos significantly wider distributions at most time points in the antibiotic-sensitive 19F (**D**) and T4 (**E**) and nutrient-sensitive D39 (**F**) comparing to respective antibiotic/nutrient-adapted or nutrient-independent strains. Vancomycin: green; Levofloxacin: blue; Kanamycin: yellow; Rifampicin: pink. For **D-F** ***: p<0.0001, **: 0.0001<p<0.001, *: 0.001<p<0.05 in a Kolmogorov-Smirnov test.

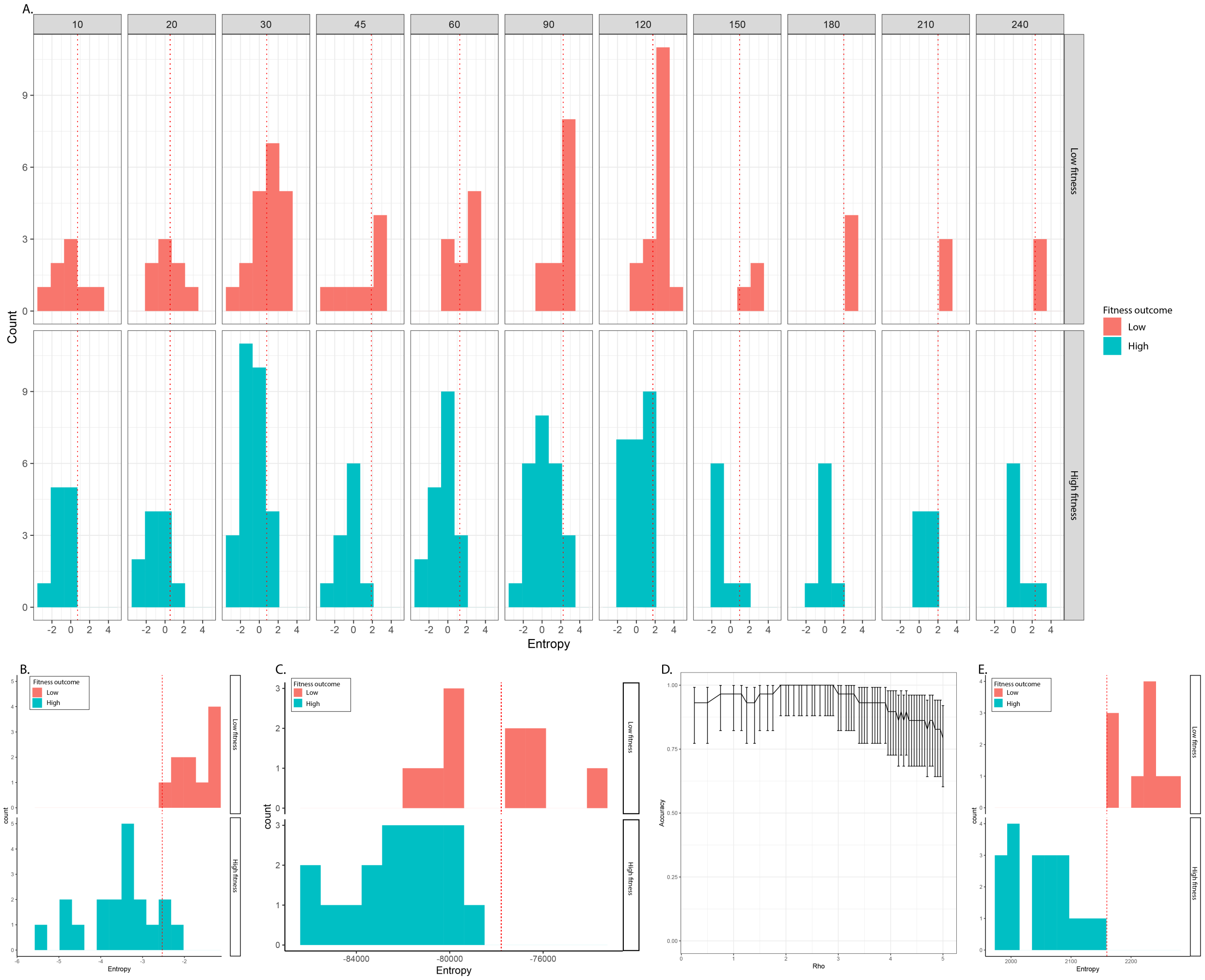


**Supplemental Figure 3.** Performance of single time point (**A**) and temporal (**B-E**) entropy prediction models on *S. pneumoniae*.

Fitness prediction can be made using timepoint-specific thresholds (red dashed line), i.e. below threshold is predicted as high fitness and above threshold is predicted as low fitness. Performance of temporal models 1, 2, 3 are shown in **B**, **C** and **E**, respectively. **D.** Accuracy as a function of regularization strength (Rho, ρ) for temporal model 3. Error bars: 95% confidence interval.

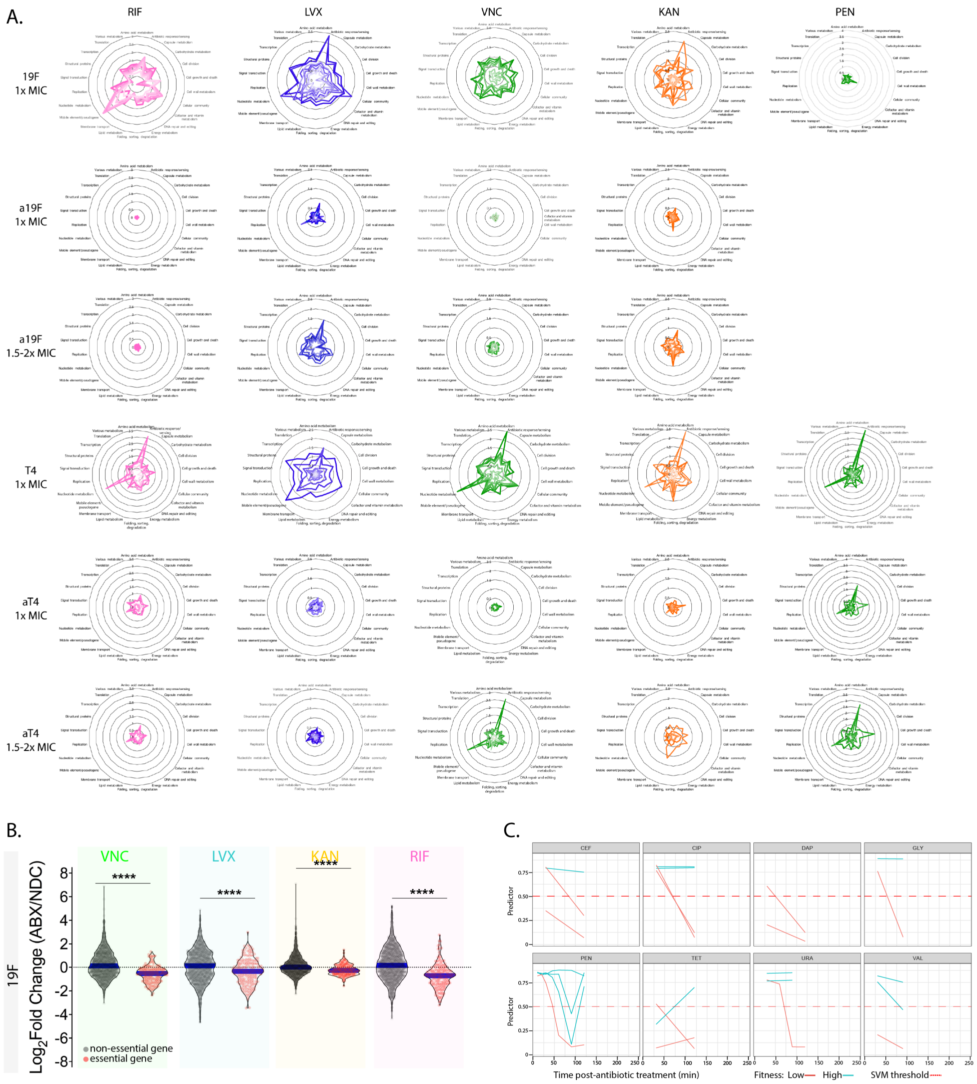


**Supplemental Figure 4.** Complex feature SVM classifier for fitness prediction.

**A.** Antibiotic triggered transcriptomic disruption can be represented by temporal entropy change at a cellular system level. Entropy in most cellular systems tend to increase over time in sensitive strains but remains unchanged in adapted strains. Transcriptional entropy is calculated for each cellular system at each time point and represented as radar plots for each temporal RNA-Seq experiment. Within a radar plot, light to dark colors indicate early (10min) to late (up to 240min) time points post-antibiotic exposure. Pink: RNA synthesis inhibitor (RIF), blue: DNA synthesis inhibitor (LVX), green: cell wall synthesis inhibitor (VNC, PEN) and yellow: protein synthesis inhibitor (KAN). T4/19F: wild-type T4/19F; aT4/19F: adapted T4/19F. **B.** When the sensitive 19F is exposed to antibiotics, essential genes are significantly more down-regulated than up-regulated comparing to non-essential genes (unpaired t-test, *: 0.001<p<0.05; **: 0.0001<p<0.001; ***: p<0.0001). **C.** Prediction probabilities generated by the CFC is plotted at each time point for strains with high fitness (cyan line) and low fitness (red line) in the presence of CEF, CIP, DAP, PEN, TET or GLY, URA, VAL depletion. A probability higher than the threshold (0.50; red dotted line) is predicted as high fitness; while a predictor lower than the threshold is predicted as low fitness.

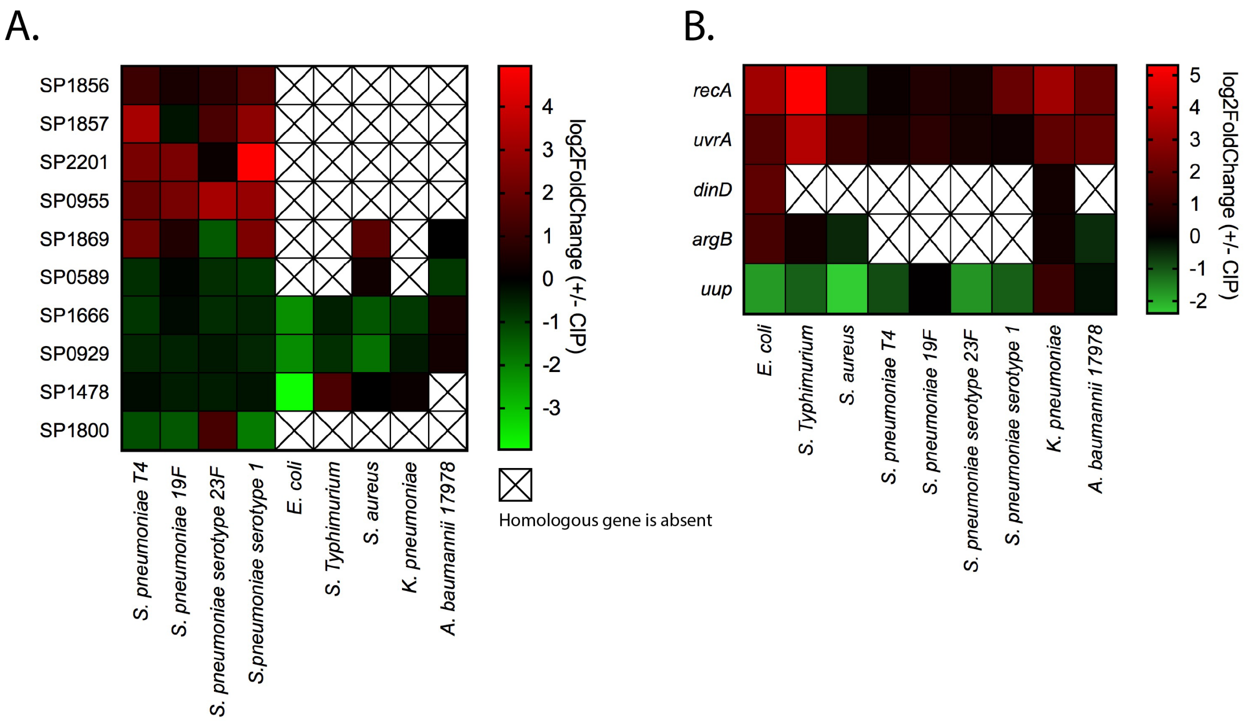


**Supplemental Figure 5.** Genes in the fitness prediction panel are poorly conserved and do not always respond in the same way among other species used in this study.

Presence and absence of each gene in the *S. pneumoniae* fitness panel (**A**.) or *E. coli* ciprofloxacin sensitivity panel (**B**; (1)) is first determined by protein BLAST based on three criteria: query coverage > 50%, E value < 1E-50 and percent identity > 30%. Differential expression of homologous genes to the *S. pneumoniae* fitness panel (**A**.) or *E. coli* ciprofloxacin sensitivity panel (**B**; (1)) do not always follow the same pattern in all six species in the presence of ciprofloxacin. For instance, *recA* and *uvrA* are not differentially expressed in *S. aureus* or *S. pneumoniae.*

**Supplemental Table 1.** Project setup

| Experimental set-up | | | | Data Collection | | Prediction | | | | |
|---|---|---|---|---|---|---|---|---|---|---|
| | | | | | | | Fitness | | | |
| Stress | Organism | Strain | Fitness outcome | Type | RNA-Seq - time points (min) | MOA | GP | $H_{temporal}$ | $H_{stp}$ | CFC |
| Vancomycin | *Streptococcus pneumoniae* | T4 | Low fitness | RNA-Seq, Tn-Seq | Temporal - 10, 20, 30, 45, 60, 90 | Train set | Train set | Train set | Train set | Train set |
| Vancomycin | *Streptococcus pneumoniae* | 19F | Low fitness | RNA-Seq, Tn-Seq | Temporal - 10, 20, 30, 45, 60, 90, 120 | Train set | Train set | Train set | Train set | Train set |
| Vancomycin | *Streptococcus pneumoniae* | aT4-VNC | High fitness | RNA-Seq, Adaptive Evolution | Temporal - 10, 20, 30, 45, 60, 90, 120 | | Train set | Train set | Train set | Train set |
| Vancomycin | *Streptococcus pneumoniae* | a19F-VNC | High fitness | RNA-Seq, Adaptive Evolution | Temporal - 10, 20, 30, 45, 60, 90, 120 | | Train set | Train set | Train set | Train set |
| Penicillin | *Streptococcus pneumoniae* | T4 | Low fitness | RNA-Seq, Tn-Seq | Temporal - 10, 20, 30, 45, 60, 90, 120 | Train set | Train set | Train set | Train set | Train set |
| Penicillin | *Streptococcus pneumoniae* | 19F | Low fitness | RNA-Seq, Tn-Seq | Temporal - 10, 20, 30, 45, 60, 90, 120 | Train set | Train set | Train set | Train set | Train set |
| Penicillin | *Streptococcus pneumoniae* | aT4-PEN | High fitness | RNA-Seq, Adaptive Evolution | Temporal - 10, 20, 30, 45, 60, 90, 120 | | Train set | Train set | Train set | Train set |
| Cefepime | *Streptococcus pneumoniae* | T4 | Low fitness | RNA-Seq, Tn-Seq | 2 time points - 30, 120 | Test set | Test set | | Test set | Test set |
| Cefepime | *Streptococcus pneumoniae* | 19F | High fitness, Low fitness | RNA-Seq, Tn-Seq | 2 time points - 30, 120 | Test set | Test set | | Test set | Test set |
| Daptomycin | *Streptococcus pneumoniae* | T4 | Low fitness | RNA-Seq, Tn-Seq | 2 time points - 30, 120 | | Test set | | Test set | Test set |
| Daptomycin | *Streptococcus pneumoniae* | 19F | Low fitness | RNA-Seq, Tn-Seq | 2 time points - 30, 120 | | Test set | | Test set | Test set |
| Levofloxacin | *Streptococcus pneumoniae* | T4 | Low fitness | RNA-Seq, Tn-Seq | Temporal - 10, 20, 30, 45, 60, 90, 120, 180 | Train set | Train set | Train set | Train set | Train set |
| Levofloxacin | *Streptococcus pneumoniae* | 19F | Low fitness | RNA-Seq, Tn-Seq | Temporal - 30, 60, 90, 120, 150, 180, 210, 240 | Train set | Train set | Train set | Train set | Train set |
| Levofloxacin | *Streptococcus pneumoniae* | aT4-LVX | High fitness | RNA-Seq, Adaptive Evolution | Temporal - 30, 60, 90, 120, 150, 180, 210, 240 | | Train set | Train set | Train set | Train set |
| Levofloxacin | *Streptococcus pneumoniae* | a19F-LVX | High fitness | RNA-Seq, Adaptive Evolution | Temporal - 30, 60, 90, 120, 150, 180, 210, 240 | | Train set | Train set | Train set | Train set |
| Ciprofloxacin | *Streptococcus pneumoniae* | T4 | Low fitness | RNA-Seq, Tn-Seq | 2 time points - 30, 120 | Test set | Test set | | Test set | Test set |
| Ciprofloxacin | *Streptococcus pneumoniae* | 19F | Low fitness | RNA-Seq, Tn-Seq | 2 time points - 30, 120 | Test set | Test set | | Test set | Test set |
| Ciprofloxacin | *Streptococcus pneumoniae* | aT4-CIP | High fitness | RNA-Seq, Adaptive Evolution | 2 time points - 30, 120 | | Test set | | Test set | Test set |
| Ciprofloxacin | *Streptococcus pneumoniae* | a19F-CIP | High fitness | RNA-Seq, Adaptive Evolution | 2 time points - 30, 120 | | Test set | | Test set | Test set |
| Kanamycin | *Streptococcus pneumoniae* | T4 | Low fitness | RNA-Seq, Tn-Seq | Temporal - 10, 20, 30, 45, 60, 90, 120, 150, 180, 210, 240 | Train set | Train set | Train set | Train set | Train set |
| Kanamycin | *Streptococcus pneumoniae* | 19F | Low fitness | RNA-Seq, Tn-Seq | Temporal - 10, 20, 30, 45, 60, 90, 120, 150, 180, 210, 240 | Train set | Train set | Train set | Train set | Train set |
| Kanamycin | *Streptococcus pneumoniae* | aT4-KAN | High fitness | RNA-Seq, Adaptive Evolution | Temporal - 30, 60, 90, 120, 150, 180, 210, 240 | | Train set | Train set | Train set | Train set |
| Kanamycin | *Streptococcus pneumoniae* | a19F-KAN | High fitness | RNA-Seq, Adaptive Evolution | Temporal - 30, 60, 90, 120, 150, 180, 210, 240 | | Train set | Train set | Train set | Train set |
| Tetracycline | *Streptococcus pneumoniae* | T4 | Low fitness | RNA-Seq, Tn-Seq | 2 time points - 30, 120 | Test set | Test set | | Test set | Test set |
| Tetracycline | *Streptococcus pneumoniae* | 19F | High fitness, Low fitness | RNA-Seq, Tn-Seq | 2 time points - 30, 120 | Test set | Test set | | Test set | Test set |
| Rifampicin | *Streptococcus pneumoniae* | T4 | Low fitness | RNA-Seq, Tn-Seq | Temporal - 10, 20, 30, 45, 60, 90, 120 | Train set | Train set | Train set | Train set | Train set |
| Rifampicin | *Streptococcus pneumoniae* | 19F | Low fitness | RNA-Seq, Tn-Seq | Temporal - 10, 20, 30, 45, 60, 90, 120 | Train set | Train set | Train set | Train set | Train set |
| Rifampicin | *Streptococcus pneumoniae* | aT4-RIF | High fitness | RNA-Seq, Adaptive Evolution | Temporal - 10, 20, 30, 45, 60, 90, 120 | | Train set | Train set | Train set | Train set |
| Rifampicin | *Streptococcus pneumoniae* | a19F-RIF | High fitness | RNA-Seq, Adaptive Evolution | Temporal - 10, 20, 30, 45, 60, 90, 120 | | Train set | Train set | Train set | Train set |
| Ciprofloxacin | *Streptococcus pneumoniae* | 23F | High fitness, Low fitness | RNA-Seq | 1 time point - 120 | | | | Validation set | |
| Ciprofloxacin | *Streptococcus pneumoniae* | 1 | Low fitness | RNA-Seq | 1 time point - 120 | | | | Validation set | |
| Ciprofloxacin | *Salmonella Typhimurium* | AR635 (CDC) | Low fitness | RNA-Seq | 1 time point - 120 | | | | Validation set | |
| Ciprofloxacin | *Staphylococcus aureus* | Mn6 (UCSD) | High fitness, Low fitness | RNA-Seq | 1 time point - 120 | | | | Validation set | |
| Ciprofloxacin | *Escherichia coli* | AR538 (CDC) | High fitness, Low fitness | RNA-Seq | 1 time point - 120 | | | | Validation set | |
| Ciprofloxacin | *Klebsiella pneumoniae* | AR497 (CDC) | High fitness, Low fitness | RNA-Seq | 1 time point - 120 | | | | Validation set | |
| Ciprofloxacin | *Acinetobacter baumannii* | ATCC17978 | Low fitness | RNA-Seq | 1 time point - 120 | | | | Validation set | |
| Ciprofloxacin | *Acinetobacter baumannii* | LAC-4 | High fitness | RNA-Seq | 1 time point - 120 | | | | Validation set | |
| No uracil | *Streptococcus pneumoniae* | T4 | High fitness | RNA-Seq, Tn-Seq | 2 time points - 30, 120 | | Train set | Train set | Train set | Train set |
| No uracil | *Streptococcus pneumoniae* | D39 | Low fitness | RNA-Seq | 4 time points - 30, 60, 90, 120 | | Train set | Train set | Train set | Train set |
| No uracil | *Streptococcus pneumoniae* | aD39-URA | High fitness | RNA-Seq, Adaptive Evolution | 2 time points - 30, 120 | | Train set | Train set | Train set | Train set |
| No L-Valine | *Streptococcus pneumoniae* | T4 | High fitness | RNA-Seq, Tn-Seq | 2 time points - 30, 120 | | Train set | Train set | Train set | Train set |
| No L-Valine | *Streptococcus pneumoniae* | D39 | Low fitness | RNA-Seq | 2 time points - 30, 120 | | Train set | Train set | Train set | Train set |
| No L-Valine | *Streptococcus pneumoniae* | aD39-VAL | High fitness | RNA-Seq, Adaptive Evolution | 2 time points - 30, 120 | | Train set | Train set | Train set | Train set |
| No Glycine | *Streptococcus pneumoniae* | T4 | High fitness | RNA-Seq, Tn-Seq | 2 time points - 30, 120 | | Train set | Train set | Train set | Train set |
| No Glycine | *Streptococcus pneumoniae* | D39 | Low fitness | RNA-Seq | 2 time points - 30, 120 | | Train set | Train set | Train set | Train set |

Cell wall synthesis inhibitor
Membrane disruptor
DNA synthesis inhibitor
Protein synthesis inhibitor
RNA synthesis inhibitor
Single nutrient depletion

High fitness
Low fitness

RNA-Seq
Tn-Seq
Adaptive Evolution

Train set
Test set
Validation set

MOA: mechanism of action
GP: gene panel
$H_{temporal}$: temporal entropy models
$H_{stp}$: single time point entropy model
CFC: complex features classifier

**Supplemental Table 2**. Genes in MOA and Fitness prediction gene-panels

| Gene panel | T4 Locus tag | Name | Product | Functional Category | Functional Tag |
|---|---|---|---|---|---|
| MOA | SP_0021 | *dut* | deoxyuridine 5'-triphosphate nucleotidohydrolase | Nucleotide metabolism | METABOLISM |
| MOA | SP_0032 | | DNA polymerase I | Replication | GENETIC INFORMATION PROCESSING |
| MOA | SP_2229 | | tryptophanyl-tRNA synthetase | Translation | GENETIC INFORMATION PROCESSING |
| MOA | SP_1073 | *rpoD* | RNA polymerase sigma factor | Transcription | GENETIC INFORMATION PROCESSING |
| MOA | SP_0338 | | ATP-dependent Clp protease ATP-binding subunit | Folding, sorting, degradation | GENETIC INFORMATION PROCESSING |
| MOA | SP_2141 | | glycosyl hydrolase-like protein | Cell wall metabolism | METABOLISM |
| MOA | SP_0798 | *ciaR* | DNA-binding response regulator CiaR | Signal transduction; Transcription | ENVIRONMENTAL INFORMATION PROCESSING; GENETIC INFORMATION PROCESSING |
| MOA | SP_2107 | | 4-alpha-glucanotransferase | Carbohydrate metabolism | METABOLISM |
| Fitness | SP_1857 | | cation efflux system protein | Membrane transport | ENVIRONMENTAL INFORMATION PROCESSING |
| Fitness | SP_1856 | | MerR family transcriptional regulator | Transcription | GENETIC INFORMATION PROCESSING |
| Fitness | SP_2201 | | choline binding protein D | Cellular community | CELLULAR PROCESSES |
| Fitness | SP_0955 | *celB* | competence protein CelB | Cellular community | CELLULAR PROCESSES |
| Fitness | SP_1869 | | iron ABC transporter permease | Membrane transport | ENVIRONMENTAL INFORMATION PROCESSING |
| Fitness | SP_0589 | | serine acetyltransferase | Amino acid metabolism | METABOLISM |
| Fitness | SP_1666 | *ftsZ* | cell division protein FtsZ | Cell division | CELLULAR PROCESSES |
| Fitness | SP_0929 | | ribosomal large subunit pseudouridine synthase D | Translation | GENETIC INFORMATION PROCESSING |
| Fitness | SP_1478 | | aldo/keto reductase | Various metabolism | METABOLISM |
| Fitness | SP_1800 | | transcriptional activator | Transcription | GENETIC INFORMATION PROCESSING |

**Supplemental Table 3**. Log2Fold changes of genes in MOA gene-panel and prediction scores (Probability) of all datasets.

| Data | SP_0021 | SP_0032 | SP_2229 | SP_1073 | SP_0338 | SP_2141 | SP_0798 | SP_2107 | Group | MOA | PredictedClass | Probability |
|---|---|---|---|---|---|---|---|---|---|---|---|---|
| T4_KAN-10 | 0.073 | 0.074 | -0.060 | -0.074 | -0.170 | -0.191 | -0.248 | -0.303 | Train | PSI | PSI | 0.118 |
| T4_KAN-20 | -0.111 | -0.066 | 0.263 | -0.144 | -0.248 | -0.418 | -0.401 | -0.609 | Train | PSI | PSI | 0.101 |
| T4_KAN-30 | -0.047 | -0.046 | -0.176 | -0.061 | 1.426 | -0.113 | -0.099 | 0.223 | Train | PSI | PSI | 0.069 |
| T4_KAN-45 | 0.033 | -0.153 | 0.082 | -0.016 | 2.560 | -0.134 | -0.105 | 0.137 | Train | PSI | PSI | 0.082 |
| T4_KAN-60 | -0.156 | -0.064 | -0.278 | -0.010 | 0.098 | -0.202 | 0.018 | 0.059 | Train | PSI | PSI | 0.064 |
| T4_KAN-90 | 0.260 | -0.232 | -0.018 | -0.136 | 0.912 | -0.115 | 0.153 | -0.079 | Train | PSI | PSI | 0.115 |
| T4_KAN-120 | -0.156 | -0.125 | -0.332 | -0.148 | 3.267 | 0.005 | -0.154 | 0.075 | Train | PSI | PSI | 0.057 |
| T4_KAN-150 | -0.177 | -0.243 | -0.635 | -0.107 | 4.961 | -0.020 | -0.304 | 0.325 | Train | PSI | PSI | 0.060 |
| T4_KAN-180 | -0.169 | -0.638 | -0.581 | 0.214 | 6.303 | -0.363 | 0.333 | 0.758 | Train | PSI | PSI | 0.065 |
| T4_KAN-210 | 0.013 | -0.797 | -1.002 | 0.245 | 6.706 | 0.179 | 1.204 | 1.449 | Train | PSI | PSI | 0.082 |
| T4_KAN-240 | -0.018 | -0.802 | -1.255 | 0.275 | 8.089 | -0.656 | 0.770 | 1.232 | Train | PSI | PSI | 0.083 |
| T4_LVX-10 | 0.008 | -0.066 | -0.095 | -0.053 | 0.053 | -0.103 | -0.038 | -0.115 | Train | DSI | PSI | 0.097 |
| T4_LVX-20 | 0.238 | 0.061 | 0.286 | 0.106 | 0.116 | 0.021 | 0.123 | 0.021 | Train | DSI | PSI | 0.318 |
| T4_LVX-30 | 0.425 | 0.246 | 0.246 | 0.027 | 0.105 | -0.029 | 0.004 | 0.018 | Train | DSI | PSI | 0.446 |
| T4_LVX-45 | 0.796 | 0.440 | 0.817 | -0.063 | -0.085 | 0.068 | 0.104 | 0.208 | Train | DSI | DSI | 0.848 |
| T4_LVX-60 | 0.959 | 0.683 | 0.698 | -0.192 | 0.068 | 0.403 | -0.031 | 0.419 | Train | DSI | DSI | 0.891 |
| T4_LVX-90 | 1.601 | 1.102 | 1.396 | -0.119 | 0.226 | 0.654 | 0.172 | 0.998 | Train | DSI | DSI | 0.925 |
| T4_LVX-120 | 2.122 | 1.678 | 1.793 | -0.428 | 0.851 | 1.127 | 0.148 | 1.677 | Train | DSI | DSI | 0.895 |
| T4_LVX-180 | 2.668 | 2.084 | 1.752 | -0.900 | 2.359 | 1.667 | 0.033 | 1.657 | Train | DSI | DSI | 0.869 |
| T4_PEN-10 | 0.154 | -0.125 | 0.110 | -0.158 | -0.141 | 0.019 | -0.181 | -0.056 | Train | CWSI | PSI | 0.175 |
| T4_PEN-20 | 0.160 | -0.125 | 0.063 | -0.160 | -0.258 | -0.032 | 0.270 | 0.350 | Train | CWSI | PSI | 0.192 |
| T4_PEN-30 | -0.009 | -0.170 | -0.064 | -0.188 | -0.187 | -0.060 | 0.970 | 0.980 | Train | CWSI | CWSI | 0.071 |
| T4_PEN-45 | 0.034 | -0.117 | -0.260 | -0.320 | -0.144 | 0.014 | 1.133 | 1.954 | Train | CWSI | CWSI | 0.038 |
| T4_PEN-60 | -0.239 | -0.160 | -0.526 | -0.503 | 0.207 | 0.036 | 1.760 | 2.684 | Train | CWSI | CWSI | 0.030 |
| T4_PEN-90 | -0.522 | -0.142 | -0.842 | -0.262 | 1.349 | 0.493 | 1.954 | 2.655 | Train | CWSI | CWSI | 0.030 |
| T4_PEN-120 | -0.220 | -0.395 | -0.666 | 0.255 | 1.691 | 1.078 | 1.778 | 2.830 | Train | CWSI | CWSI | 0.044 |
| T4_RIF-10 | -0.298 | -0.269 | -0.179 | 0.656 | 0.326 | 0.756 | -0.631 | -0.087 | Train | RSI | RSI | 0.098 |
| T4_RIF-20 | -0.818 | -0.386 | -0.614 | 1.004 | 1.149 | 1.000 | -0.466 | 0.201 | Train | RSI | RSI | 0.025 |
| T4_RIF-30 | -0.886 | -0.399 | -0.758 | 1.164 | 1.414 | 1.382 | -0.541 | 0.099 | Train | RSI | RSI | 0.017 |
| T4_RIF-45 | -1.019 | -0.589 | -0.644 | 1.724 | 1.403 | 1.014 | -0.454 | 0.592 | Train | RSI | RSI | 0.011 |
| T4_RIF-60 | -0.926 | -0.547 | -0.441 | 1.987 | 1.327 | 0.571 | -0.277 | 0.766 | Train | RSI | RSI | 0.014 |
| T4_RIF-90 | -1.182 | -0.385 | -0.556 | 2.035 | 0.819 | 0.083 | -0.020 | 1.126 | Train | RSI | RSI | 0.023 |
| T4_RIF-120 | -1.191 | -0.612 | -0.243 | 2.310 | 0.710 | 0.243 | 0.319 | 1.915 | Train | RSI | RSI | 0.029 |
| T4_VNC-10 | -0.230 | 0.018 | -0.377 | -0.252 | 0.022 | 0.179 | 0.699 | 0.936 | Train | CWSI | CWSI | 0.055 |
| T4_VNC-20 | -0.530 | -0.153 | -0.407 | -0.718 | 0.047 | 0.329 | 1.379 | 2.051 | Train | CWSI | CWSI | 0.022 |
| T4_VNC-30 | -0.742 | -0.275 | -0.695 | -1.329 | 0.233 | 0.627 | 1.621 | 2.790 | Train | CWSI | CWSI | 0.035 |
| T4_VNC-45 | -0.755 | -0.331 | -1.063 | -1.326 | 0.581 | 0.892 | 1.872 | 3.067 | Train | CWSI | CWSI | 0.043 |
| T4_VNC-60 | -0.778 | -0.457 | -0.856 | -1.127 | 0.755 | 0.859 | 2.203 | 3.304 | Train | CWSI | CWSI | 0.046 |
| T4_VNC-90 | -0.619 | -0.478 | -0.843 | -0.534 | 1.037 | 0.705 | 2.565 | 3.792 | Train | CWSI | CWSI | 0.060 |
| 19F_KAN-10 | -0.027 | -0.079 | -0.082 | 0.043 | -0.024 | -0.056 | -0.058 | -0.043 | Train | PSI | PSI | 0.099 |
| 19F_KAN-20 | -0.042 | 0.101 | -0.060 | 0.188 | 0.111 | 0.310 | -0.085 | 0.219 | Train | PSI | PSI | 0.151 |
| 19F_KAN-30 | -0.051 | -0.047 | -0.101 | 0.053 | 0.040 | 0.037 | 0.013 | -0.019 | Train | PSI | PSI | 0.102 |
| 19F_KAN-45 | -0.098 | -0.006 | -0.009 | -0.022 | 0.139 | -0.003 | -0.103 | 0.041 | Train | PSI | PSI | 0.107 |
| 19F_KAN-60 | -0.135 | -0.242 | -0.123 | -0.002 | 0.545 | -0.446 | 0.150 | -0.146 | Train | PSI | PSI | 0.052 |
| 19F_KAN-90 | -0.082 | -0.122 | -0.045 | -0.054 | 1.777 | -0.159 | 0.033 | -0.190 | Train | PSI | PSI | 0.063 |
| 19F_KAN-120 | 0.287 | -0.224 | -0.034 | -0.054 | 2.770 | -0.265 | -0.096 | 0.123 | Train | PSI | PSI | 0.085 |
| 19F_KAN-150 | -0.066 | 0.733 | -0.910 | -0.681 | 4.464 | 0.904 | -0.195 | 0.617 | Train | PSI | PSI | 0.081 |
| 19F_KAN-180 | 0.618 | -0.388 | -0.947 | 0.473 | 5.624 | -1.773 | -0.481 | 0.312 | Train | PSI | PSI | 0.083 |
| 19F_KAN-210 | -0.120 | -0.026 | -1.276 | -0.795 | 5.971 | 0.168 | -0.231 | 0.639 | Train | PSI | PSI | 0.073 |
| 19F_KAN-240 | 0.299 | -0.517 | -0.938 | -0.241 | 7.108 | -0.392 | 0.134 | 0.305 | Train | PSI | PSI | 0.075 |
| 19F_LVX-30 | 0.560 | 0.449 | 0.732 | -0.042 | -0.096 | 0.226 | 0.204 | 0.047 | Train | DSI | DSI | 0.790 |
| 19F_LVX-60 | 0.922 | 1.487 | 0.724 | -0.489 | 0.103 | 1.130 | -0.646 | 0.297 | Train | DSI | DSI | 0.904 |
| 19F_LVX-90 | 1.123 | 0.893 | 1.463 | -0.278 | 0.160 | 0.731 | -0.163 | 0.326 | Train | DSI | DSI | 0.941 |
| 19F_LVX-120 | 1.439 | 0.954 | 1.593 | -0.222 | 0.411 | 0.575 | -0.374 | 0.212 | Train | DSI | DSI | 0.948 |
| 19F_LVX-150 | 1.784 | 1.184 | 1.793 | -0.549 | 0.821 | 1.038 | -0.418 | 0.679 | Train | DSI | DSI | 0.940 |
| 19F_LVX-180 | 1.880 | 1.084 | 1.618 | -0.512 | 1.557 | 1.209 | -0.718 | 0.677 | Train | DSI | DSI | 0.923 |
| 19F_LVX-210 | 2.476 | 0.918 | 1.404 | -0.585 | 1.640 | 1.489 | -1.088 | 1.197 | Train | DSI | DSI | 0.883 |
| 19F_LVX-240 | 2.185 | 1.606 | 1.566 | -0.874 | 1.129 | 1.906 | -1.468 | 1.081 | Train | DSI | DSI | 0.872 |

| | | | | | | | | | | | | |
|---|---|---|---|---|---|---|---|---|---|---|---|---|
| 19F_RIF-10 | -0.976 | 0.154 | -1.299 | 1.252 | 0.404 | 1.405 | -0.357 | 0.342 | Train | RSI | RSI | 0.026 |
| 19F_RIF-20 | -0.779 | 0.391 | -1.368 | 2.266 | 0.849 | 2.515 | 0.172 | 0.630 | Train | RSI | RSI | 0.027 |
| 19F_RIF-30 | -0.618 | 0.188 | -1.166 | 2.827 | 1.108 | 2.874 | 0.309 | 0.402 | Train | RSI | RSI | 0.031 |
| 19F_RIF-45 | -0.378 | -0.213 | -1.249 | 3.349 | 1.395 | 3.217 | -0.051 | 0.111 | Train | RSI | RSI | 0.041 |
| 19F_RIF-60 | -0.262 | -0.286 | -0.879 | 3.681 | 1.936 | 2.834 | 0.228 | 1.078 | Train | RSI | RSI | 0.042 |
| 19F_RIF-90 | -0.282 | -0.697 | -0.724 | 2.744 | 1.843 | 2.438 | -0.368 | 1.084 | Train | RSI | RSI | 0.020 |
| 19F_RIF-120 | -0.526 | -1.727 | -0.245 | 1.953 | 1.943 | 1.740 | -1.462 | 1.383 | Train | RSI | RSI | 0.036 |
| 19F_VNC-10 | -0.007 | 0.031 | 0.181 | -0.011 | -0.005 | -0.218 | 0.163 | 0.035 | Train | CWSI | PSI | 0.174 |
| 19F_VNC-20 | -0.512 | 0.016 | -0.041 | -0.119 | -0.108 | 0.003 | 0.349 | 0.692 | Train | CWSI | CWSI | 0.072 |
| 19F_VNC-30 | -0.482 | 0.532 | -0.460 | -0.098 | 0.254 | 0.499 | 0.959 | 1.355 | Train | CWSI | CWSI | 0.055 |
| 19F_VNC-45 | -0.596 | -0.014 | -0.691 | -0.239 | 0.029 | 0.717 | 1.448 | 1.321 | Train | CWSI | CWSI | 0.029 |
| 19F_VNC-60 | 0.013 | 0.068 | -0.678 | -0.307 | 0.829 | 1.123 | 1.706 | 2.553 | Train | CWSI | CWSI | 0.039 |
| 19F_VNC-90 | -0.169 | -0.117 | -0.615 | -1.313 | 1.284 | 1.485 | 1.144 | 2.777 | Train | CWSI | CWSI | 0.052 |
| 19F_VNC-120 | 0.194 | -0.504 | -0.772 | -0.962 | 2.265 | 1.337 | 1.137 | 2.658 | Train | CWSI | CWSI | 0.058 |
| T4_CEF-30 | -0.649 | 0.068 | -0.508 | -0.531 | -0.118 | -0.128 | 1.261 | 1.925 | Test | CWSI | CWSI | 0.023 |
| T4_CEF-120 | -1.067 | -0.897 | -0.540 | -0.850 | 3.912 | 0.490 | 2.458 | 3.386 | Test | CWSI | CWSI | 0.150 |
| T4_CIP-30 | 0.268 | 0.282 | 0.291 | -0.099 | 0.005 | 0.308 | -0.359 | -0.170 | Test | DSI | PSI | 0.411 |
| T4_CIP-120 | 2.300 | 1.732 | 2.114 | -0.286 | 1.609 | 0.771 | 0.366 | 1.606 | Test | DSI | DSI | 0.870 |
| T4_TET-30 | -0.567 | -0.408 | -1.716 | -0.663 | 0.879 | 0.740 | -0.628 | -0.862 | Test | PSI | PSI | 0.063 |
| T4_TET-120 | -0.189 | -2.199 | -1.937 | 0.323 | 3.384 | -1.147 | 0.006 | 0.230 | Test | PSI | PSI | 0.120 |
| 19F_CEF22-30 | -0.030 | -0.016 | -0.053 | -0.054 | -0.113 | -0.005 | 0.431 | 0.546 | Test | CWSI | CWSI | 0.133 |
| 19F_CEF22-120 | -0.669 | -0.063 | -0.534 | -0.203 | 0.305 | -0.225 | 0.913 | 1.737 | Test | CWSI | CWSI | 0.023 |
| 19F_CIP-30 | 0.136 | 0.249 | 0.215 | -0.013 | -0.202 | -0.047 | -0.109 | 0.048 | Test | DSI | PSI | 0.303 |
| 19F_CIP-120 | 1.171 | 1.167 | 1.121 | -0.428 | 0.066 | 1.238 | -0.832 | 0.680 | Test | DSI | DSI | 0.910 |
| 19F_TET22-30 | -0.188 | -0.426 | 0.122 | -0.438 | -0.119 | -0.360 | -0.472 | -0.017 | Test | PSI | PSI | 0.070 |
| 19F_TET22-120 | -0.867 | -1.346 | -0.687 | 0.143 | 0.976 | 0.568 | -2.209 | -1.245 | Test | PSI | RSI | 0.091 |

**Supplemental Table 4**. Gene panel fitness prediction

| Strains | ABX | MOA | Concentration | Time(min) | Group | Fitness-Actual | Fitness-Predictor | Fitness-Prediction |
|---|---|---|---|---|---|---|---|---|
| T4 | CEF | CWSI | 1xMIC | 120 | Test | 0 | -0.0986 | 0 |
| 19F | CEF | CWSI | 1xMIC | 120 | Test | 1 | 1.2872 | 1 |
| 19F | CEF | CWSI | 2.75xMIC** | 120 | Test | 0 | 1.2483 | 1 |
| T4 | CIP | DSI | 1xMIC | 120 | Test | 0 | -0.6035 | 0 |
| aT4-CIP | CIP | DSI | 1xMIC | 120 | Test | 1 | 1.0671 | 1 |
| 19F | CIP | DSI | 1xMIC | 120 | Test | 0 | 0.4636 | 0 |
| a19F-CIP | CIP | DSI | 1xMIC | 120 | Test | 1 | 1.0419 | 1 |
| T4 | DAP | LIP | 1xMIC | 120 | Test | 0 | 0.1392 | 0 |
| 19F | DAP | LIP | 1xMIC | 120 | Test | 0 | 0.5660 | 1 |
| T4 | GLY | NTR | NA | 90 | Train | 1 | 0.8006 | 1 |
| D39 | GLY | NTR | NA | 90 | Train | 0 | 0.5598 | 1 |
| T4 | KAN | PSI | 1xMIC | 60 | Train | 0 | 0.5165 | 1 |
| T4 | KAN | PSI | 1xMIC | 90 | Train | 0 | -0.0958 | 0 |
| T4 | KAN | PSI | 1xMIC | 120 | Train | 0 | -0.0971 | 0 |
| T4 | KAN | PSI | 1xMIC | 150 | Train | 0 | -0.1373 | 0 |
| T4 | KAN | PSI | 1xMIC | 180 | Train | 0 | 0.1581 | 0 |
| T4 | KAN | PSI | 1xMIC | 210 | Train | 0 | 0.0049 | 0 |
| T4 | KAN | PSI | 1xMIC | 240 | Train | 0 | -0.2575 | 0 |
| aT4-KAN | KAN | PSI | 1xMIC | 60 | Train | 1 | 0.8649 | 1 |
| aT4-KAN | KAN | PSI | 1xMIC | 90 | Train | 1 | 1.5266 | 1 |
| aT4-KAN | KAN | PSI | 1xMIC | 120 | Train | 1 | 1.2792 | 1 |
| aT4-KAN | KAN | PSI | 1xMIC | 150 | Train | 1 | 0.7386 | 1 |
| aT4-KAN | KAN | PSI | 1xMIC | 180 | Train | 1 | 0.8956 | 1 |
| aT4-KAN | KAN | PSI | 1xMIC | 210 | Train | 1 | 0.9281 | 1 |
| aT4-KAN | KAN | PSI | 1xMIC | 240 | Train | 1 | 1.3548 | 1 |
| aT4-KAN | KAN | PSI | 2xMIC | 60 | Train | 1 | 1.3020 | 1 |
| aT4-KAN | KAN | PSI | 2xMIC | 90 | Train | 1 | 1.4783 | 1 |
| aT4-KAN | KAN | PSI | 2xMIC | 120 | Train | 1 | 1.0442 | 1 |
| aT4-KAN | KAN | PSI | 2xMIC | 150 | Train | 1 | 0.9295 | 1 |
| aT4-KAN | KAN | PSI | 2xMIC | 180 | Train | 1 | 1.3192 | 1 |
| aT4-KAN | KAN | PSI | 2xMIC | 210 | Train | 1 | 0.5375 | 1 |
| aT4-KAN | KAN | PSI | 2xMIC | 240 | Train | 1 | 1.3177 | 1 |
| 19F | KAN | PSI | 1xMIC | 60 | Train | 0 | 1.3697 | 1 |
| 19F | KAN | PSI | 1xMIC | 90 | Train | 0 | 0.3352 | 0 |
| 19F | KAN | PSI | 1xMIC | 120 | Train | 0 | -0.1279 | 0 |
| 19F | KAN | PSI | 1xMIC | 150 | Train | 0 | 0.2007 | 0 |
| 19F | KAN | PSI | 1xMIC | 180 | Train | 0 | 0.0924 | 0 |
| 19F | KAN | PSI | 1xMIC | 210 | Train | 0 | -0.0262 | 0 |
| 19F | KAN | PSI | 1xMIC | 240 | Train | 0 | 0.0684 | 0 |
| a19F-KAN | KAN | PSI | 1xMIC | 60 | Train | 1 | 0.4355 | 0 |
| a19F-KAN | KAN | PSI | 1xMIC | 90 | Train | 1 | 1.4213 | 1 |
| a19F-KAN | KAN | PSI | 1xMIC | 120 | Train | 1 | 1.4030 | 1 |
| a19F-KAN | KAN | PSI | 1xMIC | 150 | Train | 1 | 1.8906 | 1 |
| a19F-KAN | KAN | PSI | 1xMIC | 180 | Train | 1 | 1.3233 | 1 |
| a19F-KAN | KAN | PSI | 1xMIC | 210 | Train | 1 | 1.2375 | 1 |
| a19F-KAN | KAN | PSI | 1xMIC | 240 | Train | 1 | 1.2842 | 1 |
| a19F-KAN | KAN | PSI | 2xMIC | 60 | Train | 1 | 0.9195 | 1 |
| a19F-KAN | KAN | PSI | 2xMIC | 90 | Train | 1 | 1.1225 | 1 |
| a19F-KAN | KAN | PSI | 2xMIC | 120 | Train | 1 | 0.9592 | 1 |
| a19F-KAN | KAN | PSI | 2xMIC | 150 | Train | 1 | 1.3527 | 1 |
| a19F-KAN | KAN | PSI | 2xMIC | 180 | Train | 1 | 1.5303 | 1 |
| a19F-KAN | KAN | PSI | 2xMIC | 210 | Train | 1 | 1.4149 | 1 |
| a19F-KAN | KAN | PSI | 2xMIC | 240 | Train | 1 | 1.5510 | 1 |
| T4 | LVX | DSI | 1xMIC | 60 | Train | 0 | 0.1253 | 0 |
| T4 | LVX | DSI | 1xMIC | 90 | Train | 0 | 0.1616 | 0 |
| T4 | LVX | DSI | 1xMIC | 120 | Train | 0 | -0.1738 | 0 |
| T4 | LVX | DSI | 1xMIC | 180 | Train | 0 | -0.9192 | 0 |
| aT4-LVX | LVX | DSI | 1xMIC | 120 | Train | 1 | 1.5860 | 1 |
| aT4-LVX | LVX | DSI | 1xMIC | 150 | Train | 1 | 1.3143 | 1 |
| aT4-LVX | LVX | DSI | 1xMIC | 180 | Train | 1 | 1.3500 | 1 |
| aT4-LVX | LVX | DSI | 1xMIC | 210 | Train | 1 | 1.7234 | 1 |
| aT4-LVX | LVX | DSI | 1xMIC | 240 | Train | 1 | 1.2112 | 1 |
| aT4-LVX | LVX | DSI | 1xMIC | 60 | Train | 1 | 1.5457 | 1 |
| aT4-LVX | LVX | DSI | 1xMIC | 90 | Train | 1 | 1.5584 | 1 |
| aT4-LVX | LVX | DSI | 2xMIC | 60 | Train | 1 | 1.6053 | 1 |
| aT4-LVX | LVX | DSI | 2xMIC | 90 | Train | 1 | 1.0441 | 1 |
| aT4-LVX | LVX | DSI | 2xMIC | 120 | Train | 1 | 1.7037 | 1 |
| aT4-LVX | LVX | DSI | 2xMIC | 150 | Train | 1 | 0.7804 | 1 |
| aT4-LVX | LVX | DSI | 2xMIC | 180 | Train | 1 | 1.1615 | 1 |
| aT4-LVX | LVX | DSI | 2xMIC | 210 | Train | 1 | 1.2576 | 1 |
| aT4-LVX | LVX | DSI | 2xMIC | 240 | Train | 1 | 1.6764 | 1 |
| 19F | LVX | DSI | 1xMIC | 60 | Train | 0 | 0.2231 | 0 |
| 19F | LVX | DSI | 1xMIC | 90 | Train | 0 | 0.0488 | 0 |
| 19F | LVX | DSI | 1xMIC | 120 | Train | 0 | 0.0725 | 0 |
| 19F | LVX | DSI | 1xMIC | 150 | Train | 0 | -0.0470 | 0 |
| 19F | LVX | DSI | 1xMIC | 180 | Train | 0 | -0.2659 | 0 |
| 19F | LVX | DSI | 1xMIC | 210 | Train | 0 | 0.0187 | 0 |
| 19F | LVX | DSI | 1xMIC | 240 | Train | 0 | -0.0656 | 0 |
| a19F-LVX | LVX | DSI | 1xMIC | 60 | Train | 1 | 0.9051 | 1 |
| a19F-LVX | LVX | DSI | 1xMIC | 90 | Train | 1 | 0.8591 | 1 |
| a19F-LVX | LVX | DSI | 1xMIC | 120 | Train | 1 | 0.8019 | 1 |
| a19F-LVX | LVX | DSI | 1xMIC | 150 | Train | 1 | 1.3561 | 1 |
| a19F-LVX | LVX | DSI | 1xMIC | 180 | Train | 1 | 1.1988 | 1 |
| a19F-LVX | LVX | DSI | 1xMIC | 210 | Train | 1 | 1.5855 | 1 |
| a19F-LVX | LVX | DSI | 1xMIC | 240 | Train | 1 | 2.4605 | 1 |

| | | | | | | | | |
|---|---|---|---|---|---|---|---|---|
| a19F-LVX | LVX | DSI | 1.5xMIC | 60 | Train | 1 | 1.2931 | 1 |
| a19F-LVX | LVX | DSI | 1.5xMIC | 90 | Train | 1 | 1.1916 | 1 |
| a19F-LVX | LVX | DSI | 1.5xMIC | 120 | Train | 1 | 0.9788 | 1 |
| a19F-LVX | LVX | DSI | 1.5xMIC | 150 | Train | 1 | 1.1371 | 1 |
| a19F-LVX | LVX | DSI | 1.5xMIC | 180 | Train | 1 | 1.1947 | 1 |
| a19F-LVX | LVX | DSI | 1.5xMIC | 210 | Train | 1 | 1.2598 | 1 |
| a19F-LVX | LVX | DSI | 1.5xMIC | 240 | Train | 1 | 1.6282 | 1 |
| T4 | PEN | CWSI | 1xMIC | 60 | Train | 0 | 0.7498 | 1 |
| T4 | PEN | CWSI | 1xMIC | 90 | Train | 0 | 0.5045 | 1 |
| T4 | PEN | CWSI | 1xMIC | 120 | Train | 0 | -0.0743 | 0 |
| aT4-PEN | PEN | CWSI | 1xMIC | 60 | Train | 1 | 1.4046 | 1 |
| aT4-PEN | PEN | CWSI | 1xMIC | 90 | Train | 1 | 1.5280 | 1 |
| aT4-PEN | PEN | CWSI | 1xMIC | 120 | Train | 1 | 1.1605 | 1 |
| aT4-PEN | PEN | CWSI | 1.5xMIC | 60 | Train | 1 | 1.3355 | 1 |
| aT4-PEN | PEN | CWSI | 1.5xMIC | 90 | Train | 1 | 1.6028 | 1 |
| aT4-PEN | PEN | CWSI | 1.5xMIC | 120 | Train | 1 | 1.1851 | 1 |
| 19F | PEN | CWSI | 100xMIC* | 60 | Train | 1 | 1.1106 | 1 |
| 19F | PEN | CWSI | 100xMIC | 90 | Train | 1 | 1.4358 | 1 |
| 19F | PEN | RSI | 100xMIC | 120 | Train | 1 | 1.7687 | 1 |
| T4 | RIF | RSI | 1xMIC | 60 | Train | 0 | -0.6063 | 0 |
| T4 | RIF | RSI | 1xMIC | 90 | Train | 0 | 0.1160 | 0 |
| T4 | RIF | RSI | 1xMIC | 120 | Train | 0 | 0.2942 | 0 |
| aT4-RIF | RIF | RSI | 1xMIC | 60 | Train | 1 | 1.3158 | 1 |
| aT4-RIF | RIF | RSI | 1xMIC | 90 | Train | 1 | 1.3002 | 1 |
| aT4-RIF | RIF | RSI | 1xMIC | 120 | Train | 1 | 1.5598 | 1 |
| aT4-RIF | RIF | RSI | 2xMIC | 60 | Train | 1 | 1.3624 | 1 |
| aT4-RIF | RIF | RSI | 2xMIC | 90 | Train | 1 | 1.2072 | 1 |
| aT4-RIF | RIF | RSI | 2xMIC | 120 | Train | 1 | 1.6961 | 1 |
| 19F | RIF | RSI | 1xMIC | 60 | Train | 0 | -0.5966 | 0 |
| 19F | RIF | RSI | 1xMIC | 90 | Train | 0 | -0.6526 | 0 |
| 19F | RIF | RSI | 1xMIC | 120 | Train | 0 | -0.8956 | 0 |
| a19F-RIF | RIF | RSI | 1xMIC | 60 | Train | 1 | 1.2917 | 1 |
| a19F-RIF | RIF | RSI | 1xMIC | 90 | Train | 1 | 0.8674 | 1 |
| a19F-RIF | RIF | RSI | 1xMIC | 120 | Train | 1 | 1.1156 | 1 |
| a19F-RIF | RIF | RSI | 2xMIC | 60 | Train | 1 | 1.5610 | 1 |
| a19F-RIF | RIF | RSI | 2xMIC | 90 | Train | 1 | 1.9522 | 1 |
| a19F-RIF | RIF | RSI | 2xMIC | 120 | Train | 1 | 1.5701 | 1 |
| T4 | TET | PSI | 1xMIC | 120 | Test | 0 | -1.9411 | 0 |
| 19F | TET | PSI | 1xMIC | 120 | Test | 1 | 1.0861 | 1 |
| 19F | TET | PSI | 2.75xMIC** | 120 | Test | 0 | 0.2337 | 0 |
| T4 | URA | NTR | NA | 90 | Train | 1 | 1.7051 | 1 |
| D39 | URA | NTR | NA | 60 | Train | 0 | 1.1486 | 1 |
| D39 | URA | NTR | NA | 90 | Train | 0 | 0.2607 | 0 |
| D39 | URA | NTR | NA | 120 | Train | 0 | 0.5422 | 1 |
| aD39-URA | URA | NTR | NA | 90 | Train | 1 | 1.2746 | 1 |
| T4 | VAL | NTR | NA | 90 | Train | 1 | 0.3862 | 0 |
| D39 | VAL | NTR | NA | 90 | Train | 0 | -0.0851 | 0 |
| aD39-VAL | VAL | NTR | NA | 90 | Train | 1 | 0.5174 | 1 |
| T4 | VNC | CWSI | 1xMIC | 60 | Train | 0 | -0.2534 | 0 |
| T4 | VNC | CWSI | 1xMIC | 90 | Train | 0 | -0.8030 | 0 |
| aT4-VNC | VNC | CWSI | 1xMIC | 60 | Train | 1 | 1.5422 | 1 |
| aT4-VNC | VNC | CWSI | 1xMIC | 90 | Train | 1 | 1.1853 | 1 |
| aT4-VNC | VNC | CWSI | 1xMIC | 120 | Train | 1 | 1.8131 | 1 |
| aT4-VNC | VNC | CWSI | 1.5xMIC | 60 | Train | 1 | 0.2753 | 0 |
| aT4-VNC | VNC | CWSI | 1.5xMIC | 90 | Train | 1 | 0.0500 | 0 |
| aT4-VNC | VNC | CWSI | 1.5xMIC | 120 | Train | 1 | 0.4912 | 0 |
| 19F | VNC | CWSI | 1xMIC | 60 | Train | 0 | -0.5440 | 0 |
| 19F | VNC | CWSI | 1xMIC | 90 | Train | 0 | -0.6361 | 0 |
| 19F | VNC | CWSI | 1xMIC | 120 | Train | 0 | -0.9118 | 0 |
| a19F-VNC | VNC | CWSI | 1xMIC | 60 | Train | 1 | 1.3898 | 1 |
| a19F-VNC | VNC | CWSI | 1xMIC | 90 | Train | 1 | 1.4266 | 1 |
| a19F-VNC | VNC | CWSI | 1xMIC | 120 | Train | 1 | 1.1717 | 1 |
| a19F-VNC | VNC | CWSI | 1.5xMIC | 60 | Train | 1 | 0.7740 | 1 |
| a19F-VNC | VNC | CWSI | 1.5xMIC | 90 | Train | 1 | 0.3481 | 0 |
| a19F-VNC | VNC | CWSI | 1.5xMIC | 120 | Train | 1 | 1.0424 | 1 |

**Strain:** wild-type strains are *S. pneumoniae* T4 and 19F. Adapted strains are denoted as aT4/a19F/aD39-ABX/nutrient, e.g. a19F-KAN stands for kanamycin adapted 19F.

**Concentrations** are indicated in reference of the minimum inhibitory concentration (MIC) of the corresponding wild-type strain.

For fitness outcome and prediction, 1 indicates high fitness (i.e. a strain successfully grows in its environment), and 0 indicates low fitness (i.e. a strain cannot grow in its environment).

* 19F is resistant to penicillin. A concentration of 3μg/mL is used which is equivalent to 75x of T4 MIC of penicillin.

** 19F is not sensitive to 1x of T4 MIC of cefepime or tetracycline. A 19F-specific MIC of these two antibiotics (equivalent to 2.75x T4 MIC) is used.

Correct classification

Misclassification

**Supplemental Table 5.** Performance of temporal entropy prediction models

| Strain | ABX | Concentration | Entropy.v1 | Fitness-Prediction by Entropy.v1 | Entropy.v2 | Fitness-Prediction by Entropy.v2 | Entropy.v3 | Fitness-Prediction by Entropy.v3 | Fitness-Actual |
|---|---|---|---|---|---|---|---|---|---|
| 19F | KAN | 1xMIC | -1.771 | 0 | -75912.579 | 0 | 2179.034 | 0 | 0 |
| a19F-KAN | KAN | 1xMIC | -3.544 | 1 | -80165.225 | 1 | 2011.917 | 1 | 1 |
| a19F-KAN | KAN | 2xMIC | -2.420 | 1 | -78580.609 | 1 | 2082.585 | 1 | 1 |
| aT4-KAN | KAN | 1xMIC | -4.065 | 1 | -82044.542 | 1 | 2052.743 | 1 | 1 |
| aT4-KAN | KAN | 2xMIC | -3.123 | 0 | -79658.341 | 1 | 2095.488 | 1 | 1 |
| T4 | KAN | 1xMIC | -2.381 | 0 | -79686.818 | 1 | 2175.324 | 0 | 0 |
| 19F | LVX | 1xMIC | -1.477 | 0 | -76695.853 | 0 | 2248.076 | 0 | 0 |
| a19F-LVX | LVX | 1xMIC | -3.861 | 1 | -80932.607 | 1 | 2005.990 | 1 | 1 |
| a19F-LVX | LVX | 1.5xMIC | -3.151 | 1 | -80109.192 | 1 | 2037.754 | 1 | 1 |
| aT4-LVX | LVX | 1xMIC | -3.647 | 1 | -80767.014 | 1 | 2066.009 | 1 | 1 |
| aT4-LVX | LVX | 2xMIC | -3.602 | 1 | -80657.491 | 1 | 2066.510 | 1 | 1 |
| T4 | LVX | 1xMIC | -2.090 | 0 | -79662.623 | 1 | 2237.454 | 0 | 0 |
| 19F | PEN | 1xMIC | -4.113 | 1 | -82510.732 | 1 | 2005.874 | 1 | 1 |
| aT4-PEN | PEN | 1xMIC | -3.109 | 1 | -82619.246 | 1 | 2112.857 | 1 | 1 |
| aT4-PEN | PEN | 1.5xMIC | -2.743 | 1 | -81410.575 | 1 | 2156.750 | 1 | 1 |
| T4 | PEN | 1xMIC | -2.634 | 0 | -81455.987 | 1 | 2161.957 | 0 | 0 |
| 19F | RIF | 1xMIC | -1.571 | 0 | -76930.104 | 0 | 2269.083 | 0 | 0 |
| a19F-RIF | RIF | 1xMIC | -5.626 | 1 | -85749.977 | 1 | 1979.850 | 1 | 1 |
| a19F-RIF | RIF | 2xMIC | -4.947 | 1 | -84354.588 | 1 | 1984.332 | 1 | 1 |
| aT4-RIF | RIF | 1xMIC | -3.501 | 1 | -83054.566 | 1 | 2077.198 | 1 | 1 |
| aT4-RIF | RIF | 2xMIC | -3.673 | 1 | -83306.611 | 1 | 2065.882 | 1 | 1 |
| T4 | RIF | 1xMIC | -2.036 | 0 | -80172.936 | 1 | 2226.632 | 0 | 0 |
| D39 | URA | NA | -1.514 | 0 | -73826.043 | 0 | 2225.797 | 0 | 0 |
| 19F | VNC | 1xMIC | -1.622 | 0 | -76788.843 | 0 | 2228.850 | 0 | 0 |
| T4 | VNC | 1xMIC | -2.248 | 0 | -80640.886 | 1 | 2204.681 | 0 | 0 |
| a19F-VNC | VNC | 1xMIC | -5.072 | 1 | -84705.339 | 1 | 1983.976 | 1 | 1 |
| a19F-VNC | VNC | 1.5xMIC | -3.844 | 1 | -81455.998 | 1 | 2003.424 | 1 | 1 |
| aT4-VNC | VNC | 1xMIC | -4.620 | 1 | -86106.685 | 1 | 2043.742 | 1 | 1 |
| aT4-VNC | VNC | 1.5xMIC | -2.778 | 1 | -81905.177 | 1 | 2136.104 | 1 | 1 |

**Strain:** wild-type strains are *S. pneumoniae* T4 and 19F. Antibiotic adapted strains are denoted as aT4/19F-ABX, e.g. a19F-KAN for kanamycin adapted 19F.

**Concentrations** are indicated in reference of the minimum inhibitory concentration (MIC) of the corresponding wild-type strain.

**Entropy.v1** refers to temporal entropy-based prediction Model 1 with a threshold of -2.734, i.e. strains with an entropy higher than -2.734 are predicted to have low fitness (0); strains with an entropy lower than -2.734 are predicted to have high fitness (1).

**Entropy.v2** refers to temporal entropy-based prediction Model 2 with a threshold of -77795.54

**Entropy.v3** refers to temporal entropy-based prediction Model 3 with a threshold of 2159.525

**Supplemental Table 6.** Antibiotic minimum inhibitory concentrations (MIC) used in this study

| Antibiotic | 1x MIC for T4 (μg/mL) | 1x MIC for 19F (μg/mL) |
|---|---|---|
| Cefepime (CEF) | 0.8 | 2.2 |
| Ciprofloxacin (CIP) | 1.0 | 1.0 |
| Daptomycin (DAP) | 35 | 35 |
| Kanamycin (KAN) | 90 | 90 |
| Levofloxacin (LVX) | 1.0 | 1.1 |
| Penicillin (PEN) | 0.03 | 2.25 |
| Rifampicin (RIF) | 0.035 | 0.035 |
| Tetracycline (TET) | 8 | 22 |
| Vancomycin (VNC) | 0.24 | 0.24 |

### Supplemental Notes

#### Gene panel based misclassifications of MOA predictions

Although a few misclassified samples showed up in both train and test sets, they appear to fall into two categories: 1) The wrong MOA prediction is made based on transcriptional profiles from early time points where only a few genes are differentially expressed, for instance, in the training set LVX is predicted as PSI based on T4's response at 10-30min. In the test set CIP is predicted as PSI based on T4 or 19F's response at 30min (**Supplemental Table 3**). 2) The wrong prediction was made between PSI and RSI, i.e. the TET response in 19F at 120min is predicted as RSI. The PCA trajectories in **Figure 1E** indeed suggest a similarity between the transcriptional response under RSI and PSI treatment. We reason that since RSI and PSI target closely related cellular functions - transcription and translation, respectively, they might trigger similar response in bacteria.

#### Gene panel based misclassifications of fitness predictions

Misclassified fitness outcomes in training and test datasets fall into one of the following scenarios: 1) If an adapted strain shows a growth defect at 1.5-2xMIC, high fitness might be predicted as low fitness especially at a late time point when the transcriptional response intensifies, e.g. aT4-VNC at 60, 90min and 120min and a19F-VNC at 90min in the presence of 0.36 μg/mL of VNC. 2) LVX and KAN trigger a slow transcriptional response in sensitive strains (as indicated in **Figure 1B**). In such cases, the characteristic expression changes of the gene panel are not present until later time points. Therefore, a low fitness outcome of a sensitive strain can be misclassified as high fitness based on the transcriptional profile from an early time point, e.g. 19F and T4 60min post-kanamycin, in which there are too few differentially expressed genes. 3) In a few cases, misclassification happens when differential expression of the gene panel in one fitness outcome resembles its response in the opposite outcome. Such examples include 19F's response to DAP and CEF at 120min, T4 and D39's response to single nutrient depletion.

### Supplemental Methods

#### Bacterial strains, culture media and growth curve assays

*S. pneumoniae* strain TIGR4 (T4; NC_003028.3) is a serotype 4 strain originally isolated from a Norwegian patient(2, 3), Taiwan-19F (19F; NC_012469.1) is a multi-drug resistant strain(4, 5) and D39 (NC_008533) is a commonly used serotype 2 strain originally isolated from a patient about 90 years ago(6). Strain PG1 and PG19 were isolated from adults with pneumococcal bacteremia infection and included in the Pneumococcal Bacteremia Collection Nijmegen (PBCN)(7). All *S. pneumoniae* gene numbers refer to the T4 genome. Correspondence between homologous genes among *S. pneumoniae* strains and gene function annotations are described in **Supplemental File 1**. *Escherichia coli* strain AR538, *Klebsiella pneumoniae* strain AR497 and *Salmonella enterica subsp* Typhimurium strain AR635 were clinical isolates obtained from the Center of Disease Control (CDC). *Staphylococcus aureus* strain MN6 was kindly provided by George Sakoulas (Center of Immunity, Infection & Inflammation, UCSD School of Medicine). Unless otherwise specified, *S. pneumoniae* strains were cultivated in Todd Hewitt medium with 5% yeast extract (THY) with 5μL/mL oxyrase (Oxyrase, Inc) or on sheep's blood agar plates (Northeastern Laboratories) at 37°C with 5% CO2. *A. baumannii*, *E. coli*. *K. pneumoniae*, *S. aureus* and *S.* Typhimurium were cultured in Mueller Hinton broth II (Sigma) at 37°C with 220rpm constant shaking. Tn-Seq and RNA-Seq experiments of *S. pneumoniae* under nutrient-depletion and antibiotic conditions were performed in chemically defined medium (CDM)(8) and semi-defined minimal medium (SDMM)(9), respectively. RNA-Seq experiments for *A. baumannii, S.* Typhimurium, *E. coli*. *K. pneumoniae*, and *S. aureus* were performed in Mueller Hinton broth II. Single strain growth assays were performed at least three times using 96-well plates by taking $OD_{600}$ measurements on a Tecan Infinite 200 PRO plate reader.

#### Tn-Seq experiments, sample preparation and analysis

Six independent transposon libraries were constructed in T4 and Taiwan-19F using transposon Magellan 6 as previously described(9-11). Tn-Seq experiments under single nutrient depletion conditions were performed in CDM in the presence or absence of one of the three nutrients: Glycine, uracil and L-Valine. Antibiotic Tn-Seq experiments were performed in SDMM in the presence or absence of cefepime, ciprofloxacin, daptomycin(12), kanamycin, levofloxacin, penicillin, tetracycline, rifampicin, or vancomycin. Library preparation, Illumina sequencing, data processing and fitness calculations ($W_i$; representing the growth rate) were performed as previously described(9-11, 13). Genes with a significant fitness change must satisfy three criteria: 1) Fitness of a gene must be calculated from at least three insertion mutants in both control and experimental conditions; 2) A gene must have a fitness difference greater than 15% ($|W_{Control}-W_{Experimental}|>0.15$); 3) $W_{Control}$ and $W_{Experimental}$ must significantly differ in a one sample t-test with Bonferroni correction for multiple testing.

**Temporal RNA-Seq sample collection, preparation and analysis**

In nutrient RNA-Seq experiments, T4, D39 and adapted D39 were collected at 30 and 90min after depletion of D39-essential nutrients. In the training set antibiotic RNA-Seq experiments, wild-type and adapted T4 or 19F were collected at 10, 20, 30, 45, 60, 90, 120min post-vancomycin, rifampicin or penicillin treatment. Additional time points at 150, 180, 210 and 240min were collected in levofloxacin and kanamycin experiments due to the slower transcriptional response. In the test set antibiotic RNA-Seq experiments, wild-type T4 and 19F were collected at 30 and 120min post-cefepime, ciprofloxacin, daptomycin or tetracycline treatment. Ciprofloxacin-adapted T4 and 19F were collected at 30 and 120min post-ciprofloxacin treatment. Wild-type strains were exposed to 1xMIC antibiotics; antibiotic-adapted strains were exposed to 1xMIC and 1.5-2xMIC of the respective antibiotic. Cell pellets were collected by centrifugation at 4000 rpm at 4$^{o}$C and snap frozen and stored at -80$^{o}$C until RNA isolation with the RNeasy Mini Kit (Qiagen). 400ng of total RNA from each sample was used for generating cDNA libraries following the RNAtag-Seq protocol(14) as previously described(8). PCR amplified cDNA libraries were sequenced on an Illumina NextSeq500 generating a high sequencing depth of ~7.5 million reads per sample(15). RNA-Seq data was analyzed using an in-house developed analysis pipeline. In brief, raw reads are demultiplexed by 5' and 3' indices(14), trimmed to 59 base pairs, and quality filtered (96% sequence quality>Q14). Filtered reads are mapped to the corresponding reference genomes using bowtie2 with the --very-sensitive option (-D 20 –R 3 –N 0 –L 20 –i S, 1, 0.50)(16). Mapped reads are aggregated by featureCount and differential expression is calculated with DESeq2(17, 18). In each pair-wise differential expression comparison, significant differential expression is filtered based on two criteria: |log2foldchange| > 1 and adjusted p-value (padj) <0.05. All differential expression comparisons are made between the presence and absence of the antibiotic or nutrient at the same time point.

**Experimental evolution**

D39 was used as the parental strain in nutrient-depletion evolution experiments; T4 and 19F were used as parental strains in antibiotic evolution experiments. Four replicate populations were grown in fresh CDM with a decreasing concentration of uracil or L-Val for nutrient adaptation populations, or an increasing concentration of ciprofloxacin, cefepime, levofloxacin, kanamycin, penicillin, rifampicin, or vancomycin for antibiotic adaptation populations. Four replicate populations were serial passaged in CDM or SDMM as controls to identify background adaptations in nutrient or antibiotic adaptation experiments, respectively. When populations were adapted to their nutrient or antibiotic environment, a single colony was picked from each experiment and checked for its adaptive phenotype by growth curve experiments.

**Determination of relative minimal inhibitory concentration (MIC)**

1 to 5 x $10^5$ CFU of mid-exponential bacteria in 100uL was diluted with 100uL of fresh medium with a single antibiotic to achieve a final concentration gradient of cefepime (T4: 0.008-0.8 µg/mL; 19F: 0.6-2.4 µg/mL), ciprofloxacin (*S. pneumoniae* strains: 0.125-4.0 µg/mL; other species: 0.0125-25 µg/mL), daptomycin (15-55 µg/mL), levofloxacin (0.1-2 µg/mL), kanamycin (35-250 µg/mL), penicillin (T4: 0.02-0.055 µg/mL, 19F: 1-4 µg/mL), rifampicin (0.005-0.04 µg/mL), tetracycline (T4: 4-18 µg/mL; 19F: 19-22 µg/mL), and vancomycin (0.1-0.5 µg/mL) in 96-well plates. Each concentration was tested in triplicate. Growth was monitored on a Tecan Infinite 200 PRO plate reader at 37$^o$C for 16 hours. MIC is determined as the lowest concentration that abolishes bacterial growth (**Supplemental Figure 1**).

**PCA and Trajectory clustering**

For principal component analysis (PCA), differential expression (log2fold change of +/- antibiotic comparisons) data from all 255 experimental conditions (per time point per antibiotic from all experiments excluding CIP-validation set with *A. baumannii*, *E. coli*, *K. pneumoniae*, *S.* Typhimurium, *S. aureus*, *S. pneumoniae* serotype 1 and 23F strains) were assembled in R (v3.4.3). The function "prcomp" was used for PCA. Timepoints of the same experiment were connected to form trajectories. Since not all experiments are on the exact same time scale (e.g. KAN experiments extend to 240min whereas RIF experiments cover 120min), equivalent timepoints for each experiment were determined to be $\frac{i \times t_{max}}{6}$ for $i = 1,2,\ldots,6$ and $t_{max}$ being the latest time point available for the corresponding experiment. If a timepoint did not correspond to an existing RNA-Seq data point, this time point was inferred by linear interpolation of the existing trajectories. To cluster these trajectories, a trajectory-distance metric between two trajectories $X$ and $Y$ is defined as the sum of Euclidean distances ('dist', on the principal component coordinates) $\sum_{i=1}^{6} dist(X_i, Y_i)$ of all timepoints i. All pairwise distances are computed for all pairs of trajectories included in the analysis (WT strains with low fitness, for PSI, DSI, CWSI and RSI). Kmeans clustering with k=4 is used on the pairwise distances to cluster the trajectories.

**Selection of gene panel for MOA prediction**

Differential expression (log2 fold change of drug/no drug comparison) data from 39 antibiotic conditions (including VNC, LVX, RIF, PEN, KAN) from wildtype strains (with low fitness outcome) were assembled in R (v3.4.3). Genes with incomplete data (e.g. genes unique to one strain) were omitted. The differential expression data was then scaled such that the values for each gene had mean=0 and variance=1. Feature selection was done by fitting a logistic regression model with lasso regularization (setting lambda=0.3) for each MOA (DNA synthesis inhibitor, RNA synthesis inhibitor, protein synthesis inhibitor, cell wall synthesis inhibitor) independently. A total of 8 features (i.e. genes) selected for each model were then taken together, and a multi-class SVM was trained using the selected features. The performance was evaluated on a training set (VNC, LVX, RIF, PEN, KAN) and an independent test set that includes different antibiotics (CIP, CEF, TET).

**Selection of gene panel for fitness prediction**

Data from all 255 experimental conditions were assembled and standardized as described in "Selection of gene panel for MOA prediction". The data was then split into training (KAN, LVX, PEN, RIF, VNC, URA, VAL, GLY) and test (TET, CEF, DAP, CIP) sets. A logistic regression model with lasso regularization (setting lambda=0.15) was trained using the training dataset, using the glmnet package v2.0.

**Quantifying entropy of temporal transcriptional data**

Entropy of a single-time point experiment ($H_{stp}$) is calculated using the definition of entropy on normal distributions (equation 1)

$$H_{stp} = \frac{1}{2}\ln(2\pi e\sigma^2) \quad (1)$$

Where $\sigma^2$ is the variance of the differential expression distribution. A threshold was selected by scanning a range of possible thresholds and selecting one that maximizes accuracy on the training set (same as gene panel training set). Performance is reported on this training set and test set (same as gene panel test set).

While this metric is informative, it does not take into account the temporal changes that occur in genes. For the temporal RNA-Seq experiments for which multiple time points exist, the variance of differential expression is quantified using equation 2

$$H_{temporal\,1} = \sum_i \ln\left(2\pi\sigma_i^2\right) \quad (2)$$

Where $\sigma_i^2$ is the variance in differential expression of $gene_i$ over $t$ time points. Thresholding is done similarly to $H_{stp}$.

An assumption in the previous model is that genes' variation over time are independent of one another. Genes in the same regulon are known to be co-expressed, and are examples of highly covarying genes. This means there are potential correlations between pairs of genes. In order to account for this phenomenon, the expression changes in $N$ genes over $t$ timepoints are considered to come from a multivariate normal distribution, with $N$ dimensions. This is in contrast to equation 2, where $N$ independent univariate normal distributions are considered. The entropy of a multivariate normal distribution is defined as (equation 3)

$$H_{temporal\,2} = \ln(|\Sigma|) \quad (3)$$

Where $\Sigma \in \mathbb{R}^{N \times N}$ is the covariance matrix ($\Sigma_{ij}$ is the covariance of $gene_i$ and $gene_j$), and $|\Sigma|$ denotes the determinant of $\Sigma$. Thresholding is done similarly to $H_{stp}$.

It is likely that $\Sigma$ takes into account indirect links between genes. To correct for this, regularization is applied using glasso (v.1.10), which eliminates spurious links that are potentially an artifact of such indirect covariances. Glasso applies an L1-penalty to estimate a sparse inverse covariance matrix (precision matrix). The inverse of this sparse precision matrix is used as the regularized covariance matrix $\Sigma^{\rho}$ where $\rho$ denotes the regularization strength. The higher the value of $\rho$, the sparser the matrix. Multiple values of $\rho$ are scanned between 0 and 5 ($\rho$=0 being equivalent to $H_{temporal\,2}$ and $\rho$ >5 being equivalent to $H_{temporal\,1}$). For each $\rho$, the inverse-regularized-inverse covariance entropy is computed as (equation 4)

$$H_{temporal\,3} = \ln(|\Sigma^{\rho}|) \quad (4)$$

and the appropriate threshold is selected. The accuracy of each of these models is reported (with varying $\rho$,) on a training set that includes all experiments for which >3 time points are available. The final model selected has a $\rho$, and threshold value that maximizes accuracy.

**Feature selection and model training for SVM fitness predictor**

For each experimental time point, the following features are computed: entropy of differential expression (DE) (see equation 1) for all genes; entropy of DE for genes belonging to each functional category (this results in 23 features/experiment, one for each category); number of TIGs (Transcriptionally Important Genes; those genes that change in transcription as determined by RNA-Seq); entropy of fitness change (dW); entropy of dW for genes belonging to each functional category; number of PIGs (Phenotypically Important Genes; those genes that contribute to fitness changes as determined by Tn-Seq); difference between mean DE of essential genes, and mean DE of non-essential genes; proportion of essential genes that are significantly downregulated (log2FoldChange<-1, padj<0.05); evenness of functional category distribution of TIG (in order to characterize this, we compute the Kullback-Liebler Divergence (KLD) of the TIG category distribution from the category distribution of all genes present in the genome); evenness of functional category distribution of PIGs (KLD of PIG category distribution from whole-genome category distribution); similarity of category distribution of PIGs to that of TIGs (to quantify this, we use L1-distance between the category distributions of TIGs and PIGs); mechanism of action of the stress (as a categorical variable). The categorical features (such as mechanism of action) are one-hot encoded and all data is standardized such that all scalar features have mean=0 and variance=1. The data is then split into training and test set as mentioned above. Feature selection was performed using regularized logistic regression with L1 penalty, and C=0.1 in scikit-learn v0.20.2, such that ~10% of the 54 features would remain. This resulted in 6 features being selected (**Figure 5C**), including KLD of PIGs (KLD.P), proportion of downregulated essential genes (downEss), entropy of 3 categories (Folding, sorting, degradation; Signal transduction, and Mobile element/pseudogene), and whether the MOA is PSI. A support vector classifier using the default parameters in scikit-learn v0.20.2 was trained on the training data, using these 6 features. Performance is reported on the training and test data sets
